# Robust design and validation of LAMP assays for in-field detection of three major bacterial vascular diseases of banana

**DOI:** 10.1101/2025.11.10.687727

**Authors:** Isabelle Robène, Mikel Arrieta-Salgado, Samuel Rozsasi, Diane Mostert, Véronique Maillot-Lebon, Yolande Chilin-Charles, Jos Jansen van Vuuren, Gloria Valentine Nakato, Reagan S. Kanaabi, Freddy Magdama, Estefany. Paredes Salgado, Catur Hermanto, Agus Sutanto, Janet Conie, Debbie Henry, Sheryl Le Roux, Altus Viljoen, Emmanuel Jouen, Stephane Poussier, Emmanuel Wicker, Yann Pecrix

## Abstract

Bacterial diseases of banana are becoming increasingly significant worldwide, resulting in reduced yields and higher disease management costs. The most important bacterial diseases of banana include Moko and banana blood disease (BBD), caused by *Ralstonia solanacearum* and *Ralstonia syzygii* subsp. *celebesensis*, respectively, and banana *Xanthomonas* wilt (BXW) caused by *Xanthomonas vasicola* pv. *musacearum*. Effective surveillance and disease management require point-of-care diagnostics, such as loop-mediated isothermal amplification (LAMP), for on-site operation. In this study, three LAMP assays were developed to specifically detect the bacteria responsible for Moko, BBD and BXW, directly from banana tissues, using a simplified DNA extraction protocol. The BBD - and BXW-LAMP assays demonstrated 100% specificity, yielding negative results for a broad range of non-target bacteria, including closely related species as well as pathogenic and endophytic strains associated with banana, and positive results for all the tested target strains. For Moko disease, a duplex-LAMP assay was developed to detect all strains from the four globally most relevant sequevars: IIB-3, IIB-4, IIA-6, and IIA-24. The duplex-LAMP successfully detected all target strains, except one that was shown to be non-pathogenic to Cavendish bananas. All non-target strains tested negative, with the exception of a delayed signal for one strain belonging to *Ralstonia thomasi*, not associated with banana environment (hospital strain). These results were supported by an extensive *in silico* analysis conducted on 9,668 Burkholderiaceae and 7,483 Xanthomonadaceae genomes. Detection limits ranged from 0.1 pg/µl to 1 pg/µl DNA, and from 10⁴ to 10⁵ CFU/ml on banana tissues spiked with calibrated bacterial suspensions, depending on the assay. The LAMP assays prove highly effective for detecting target pathogens in both artificially inoculated banana plants and field samples, offering a promising tool for improving disease management strategies.

## Introduction

Threats to plants by invasive pathogens are increasing worldwide as a result of globalization, human mobility, climate change, and pathogen or vector evolution [1]. Emerging and re-emerging plant pathogens have the potential to cause significant economic losses. Efficient plant disease management requires disease identification through surveillance and the availability of efficient diagnostic tests to detect the causal agent directly from plant material [2]. Inaccurate diagnosis of pathogens can have a dramatic economic impact, either by allowing the introduction of harmful organisms into previously unaffected areas, or conversely by prompting the unnecessary deployment of inappropriate and expensive disease management strategies.

Banana (*Musa* spp) is the world’s most important fruit crop in terms of production volume (more than 135 million tons globally) [3] and trade. It is grown in more than 130 countries, mainly in the tropics and subtropics, where approximately 400 million people rely on the crop as a staple food and source of income. Global banana production is however constrained by several pests and diseases that are responsible for yield losses and low productivity. This severely compromises food security and livelihoods for banana-based farming households. Bacterial diseases constitute a significant part of these threats. Among them, vascular diseases such as Moko/Bugtok disease caused by *Ralstonia solanacearum*, banana blood disease (BBD) caused by *Ralstonia syzygii* subsp. *celebesensis*, and *Xanthomonas* wilt of banana and enset (BXW), caused by *Xanthomonas vasicola* pv. *musacearum* (Xvm), are of major concern due to their high virulence and significant impact on production [4].

Bacterial strains of the *R. solanacearum* species complex (*RSSC*), responsible for bacterial wilt, are considered priority plant pathogens in many countries worldwide. *RSSC* has been subdivided into three species: *R. solanacearum*, corresponding to phylotype II strains (including Moko strains), and further subdivided into subgroups IIA and IIB, *R. pseudosolanacearum*, corresponding to phylotypes I and III; and *R. syzygii*, corresponding to phylotype IV [5,6]. Additionally, *R. syzygii* sp. nov. is further divided into three subspecies: *R. syzygii* subsp. *indonesiensis* sp. nov., which includes broad host range strains, such as tomato and potato; *R. syzygii* subsp. *syzygii* sp. nov., which causes Sumatra disease of cloves; and *R. syzygii* subsp. *celebesensis* sp. nov., which includes the strains responsible for BBD.

Moko disease has been present in numerous countries of Latin America and the Caribbean [7] since its first description in British Guiana in the middle of the 19^th^ century [8], as well as in Philippines and Malaysia [9]. In Colombia, the disease has severely affected banana and plantain production in the main growing areas, causing production losses of up to 100% [10]. Additionally, Moko disease has caused significant damage to banana and plantain production in Ecuador since its first description in 2014 [11]. According to official reports, over 3,000 hectares of banana have already been affected [12]. While detailed economic estimates of losses in banana and plantain are still limited, the continuous increase in local plantain prices suggests a sharp decline in production areas and highlights the broader economic repercussions of the disease [13].

A high diversity was described among bacterial strains causing Moko disease, which have historically been classified into four *egl* sequence-based sequevars: IIA-6, IIA-24, IIB-3, and IIB-4 [14]. All these sequevars induced wilt on banana, plantain, and *Heliconia* spp. Beside these, the pathological variant IIB-4NPB, first identified in infected *Anthurium andreanum* in Martinique, is non-pathogenic on banana, but genetically very close to the Moko lineage IIB-4 [15]. Recently, Moko disease-related strains identified in Brazil clustered within Solanaceae-associated sequevars IIA-41 and IIB-25. Additionally, the authors reported a new sequevar, IIA-53 [16].

Blood disease of banana has been prevalent in Indonesia since 1905 [17] and was recently detected in Malaysia [18]. Genetic analyses revealed that there is little diversity among strains of BBD [14,19,20]. Plantation losses can be very severe (27–80%). In Sumatra, banana blood disease has caused the loss of over 20,000 tons of bananas, with an estimated economic impact of US$1 million.

Initially described in the early 1970s in Ethiopia, *Xanthomonas* wilt of banana has become the most important and widespread disease of *Musa* spp. in East and Central Africa since 2001: Uganda, Eastern Democratic Republic of Congo, Rwanda, Tanzania, Kenya and Burundi [21]. Xvm infects all *Musa* cultivars in this region and can cause up to 100% yield loss, especially in ABB type bananas [22].

Banana-infecting Xvm was found subdivided into two main sublineages based on Whole-genome SNPs [23]. Furthermore, MLVA-19 genotyping revealed an unexpected diversity by identifying several homogeneous clusters within each of these sublineages, notably in Ethiopia [24].

Common symptoms of bacterial diseases in bananas include leaf yellowing and wilting, vascular discoloration in the pseudostem and rhizome, premature fruit ripening and dry rot of the fruit. Bacterial ooze can also be observed in the internal tissues. Relying solely on symptomatology for diagnosing these diseases is not reliable, as these general vascular symptoms can lead to misdiagnosis. Furthermore, the symptoms can vary depending on the banana cultivar. As an example, Moko disease is characterized by yellowing of leaves in medium-to large-scale Cavendish plantations (*Musa accuminata* AAA, dessert bananas), while fruit rotting is the main symptom observed in *M. Balbisiana* cultivars (BBB cooking banana) [4]. Moreover, internal pseudostem discoloration and leaf yellowing symptoms can be also confused with a fungal disease, *Fusarium* wilt.

Different molecular tools are available for the detection and diagnosis of banana bacterial pathogens, mainly PCR and real-time PCR tools. For Moko disease, a Musa multiplex-PCR [25,26], a duplex-PCR [27] and more recently a revisited Musa multiplex-PCR including primers for the new Brazilian sequevars [28], are available. For BBD, PCR and quantitative PCR assays are available [29,30]. For BXW, several PCR assays were developed and particularly five specific primers pairs designed from genes unique to XVM [31]. Most of these molecular tools are reliable and sensitive, but they require expensive laboratory equipment, high sample purity, and are relatively time-consuming. On the other hand, Point-of-Care (POC) diagnostic assays that do not require sophisticated equipment and that can be conducted on-site, are gaining popularity in the fields of human, animal and plant health. Among these POC techniques, the molecular Loop-mediated isothermal amplification (LAMP) method [32] is of particular interest because it is fast, inexpensive, simple, specific and sensitive and can be used following a very simplified nucleic acid extraction method. A LAMP assay was already developed for Xvm, but the targeted *GspD* sequence was not exclusive to Xvm, possibly leading to false positive responses [33]. A LAMP protocol with detection based on flocculation of carbon particles was recently published for BBD [34]. To date, no LAMP assay is available for specifically detecting Moko disease-inducing *Ralstonia solanacearum* subgroups.

In this study, the development of highly specific and sensitive LAMP assays for the diagnosis of Moko, BBD and BXW from symptomatic plant tissues is presented. The DNA targets were selected based on an extensive comparative genomics study. A rapid DNA preparation step was implemented before the real-time LAMP (RT-LAMP) assay. The LAMP assays were successfully evaluated on inoculated banana plants as well as in-field in different countries where the diseases are present.

## Material and methods

### Bacterial strains and DNA extraction

The 15 BXW strains, 11 BBD strains and 41 Moko strains, used in this study, along with non-target strains belonging to different lineages, species or genera, were listed, along with their origin and host, in the S1, S2 and S3 Tables respectively. Strains were stored at – 80 °C on beads in cryovials (Microbank Prolab Diagnostics) or were freeze-dried for long-term storage. Both *Xanthomonas* strains and non-*RSSC* cultures were routinely grown on yeast-peptone-glucose agar (YPGA) (yeast extract 7 g/l; peptone 7 g/l; glucose 7 g/l; agar 18 g/l; pH 7.2) at 28 °C, while *RSSC* strains were grown on modified Kelman medium including tetrazolium chloride (TZC) at 28 °C [35]. Bacterial cultures grown overnight were used for DNA extraction using a DNeasy Blood and tissues kit (Qiagen, Courtaboeuf, France).

### Selection of target DNA sequences

A bioinformatics screening of candidate Coding Sequences (CDSs) was performed using the “gene phyloprofile” tool available on the MicroScope platform of the Genoscope, France [36]. For *Xanthomonas*, eight Xvm genomes were compared to 26 non-target *Xanthomonas* genomes (other species and pathovars) in order to select CDSs conserved in all Xvm genomes that had limited or no identity to CDSs from non-target genomes present in the database (S4 Table). For Moko, 10 target genomes were compared against 13 non-target *RSSC* genomes, including strains belonging to close non-target sequevars. As no target common to all the Moko strains while absent in non-target strains could be found, the analysis was performed separately for the subgroup IIB strains (n=6) and the subgroup IIA (n=4). For BBD one target genome was compared to 24 non-target genomes, including closely related genomes belonging to the other subspecies of *Ralstonia syzygii*.

In order to screen the different candidate markers, BLASTN searches were performed against NCBI databases using the selected nucleotide sequences as queries. Searches for BBD and Moko targets were restricted to the *Ralstonia* group (taxid:119060): nr/nt, draft (n = 2,308,262 contigs), complete genomes (n = 2,653), and RefSeq representative genomes (n = 17,601). For the BXW target, searches were restricted to the *Xanthomonas* group (taxid:32033): nr/nt, draft (n = 1,802,950 contigs), complete genomes (n = 1,870), and RefSeq representative genomes (n = 8,544).

### LAMP primers design

LAMP primers were designed using Primer explorer V5 software (https://primerexplorer.eiken.co.jp/lampv5e/index.html). Several primer sets were designed for the different pathogens and experimentally screened on a set of target and non-target strains in order to select the most effective one.

### Real Time-LAMP assays

LAMP reactions were performed with a portable fluorometer device (Genie II, OptiGene, Horsham, UK), in 25 μl total reaction volume, containing 15 μl ISO-DR004 Isothermal Mastermix (OptiGene, Horsham, UK), with 2.5 μl of pre-primer mix, giving final concentrations of 0.2 μM of each F3 and B3 primers, 0.8 μM of each FIP and BIP primers, 0.4 μM of each loop primer, 2.5 μl pure H_2_O and 5 μl of template DNA. These final standard primer concentrations were referred to as “1X”. The duplex-MOKO LAMP assay was performed under similar conditions except that the mix received 2.5 μl of each of the two LAMP primer sets (1X each), and no water. Standard LAMP reactions were run at 65 °C for a period of 30 min, followed by an anneal step with temperatures varying from 98 °C to 80 °C at a speed of 0.05 °C/second. Amplification curves (associated to a “Time to Result value” = TTR values) and annealing temperature (Ta) peaks were generated during the LAMP reaction.

### Optimization of the LAMP protocols

For each LAMP assay, a temperature gradient analysis (61° C - 68° C) was performed to determine the temperature associated with the earliest Time to Result (TTR) signals. These analyses were performed at least in duplicate using simplified DNA extracts from plant material spiked with a bacterial concentration of 10^5^ CFU/ml. For the BXW LAMP system, the standard mix (1X) was compared to a mix where all the primer concentrations were doubled (2X).

### Simplified DNA extraction protocol

A quick alkaline extraction method (NaOH method) modified from [37] was used on the three bacterial pathogens for DNA preparation from infected plant material. About 100 mg of healthy pseudostem tissue was macerated for 10 min in 1 ml of freshly prepared 0.5 M NaOH including 2% Polyvinylpyrrolidone (PVP40) (Sigma-Aldrich, Saint-Quentin Fallavier, France). Five microliters of the homogenate were further diluted with 195 μl 100 mM Tris (pH 8.0). This homogenate was then used immediately for the LAMP assay with the ISO-DR004 Isothermal Mastermix.

### Specificity

#### In silico analysis

*In silico* specificity of the Moko and BBD-specific candidates was assessed on Burkholderiaceae genomes, while BXW-specific candidates were tested against Xanthomonadaceae genomes. All publicly available genome assemblies belonging to Burkholderiaceae and Xanthomonadaceae were obtained from the NCBI GenBank database on October 24th and November 29th, 2024, respectively. Additionally, for the Burkholderiaceae *in silico* analysis, 336 unpublished genome assemblies available in-house were included. All assemblies were filtered using BUSCO (version 5.5.0) for their completeness (>95%) and duplication level (≤10%). We retained 9,668 genomes for the Burkholderiaceae and 7,483 genomes for the Xanthomonadaceae. Homologous sequences were identified by BLASTN (version 2.9.0) and extracted using BEDTools getfasta (version 2-2.30.0). For each of the genomic regions targeted by the different LAMP systems, a list of unique allelic variants was defined by removing redundant sequences using SeqKit rmdup (version 2.8.1) and aligned using MUSCLE (version 5.1.0). Full -length targets and LAMP primers specificity were defined from these alignments. The percentage identity of each primer to its target sequence was calculated using BLASTN.

### Experimental data

Analytical specificity was evaluated on DNA of target and non-target strains adjusted to 1 ng/µl, following the guidelines in the EPPO PM 7/98 standard and using two criteria: inclusivity, i.e. the ability of the LAMP systems to detect all strains of the target organism; and exclusivity, i.e. the capacity to generate negative results from non-target strains [38].

### Sensitivity

The sensitivity of the real-time LAMP assays was assayed on (i) serial dilutions of bacterial DNA (0.1 ng/µl to 0.0001 ng/µl) extracted with the DNeasy Blood Tissue kit (Qiagen, Courteboeuf, France) according to the supplier’s instructions and tested in quadruplicate by the dedicated LAMP assays; and (ii) calibrated bacterial suspensions added to banana tissues, from 1 × 10^6^ CFU /ml to 1 × 10^3^ CFU/ml, eight replicates for 1 × 10^6^ CFU /ml, 10-12 replicates for other dilutions. For (ii), bacterial suspensions from overnight cultures on modified Kelman or on YPGA plates (Xanthomonads strains) were adjusted spectrophotometrically to a concentration of 1 × 10^8^ CFU /ml and serially diluted in 0.01 M Tris buffer pH 7.2. Bacterial cell concentrations were checked by plating 50 μl of the 10^4^ dilution on modified Kelman or on YPGA plates (Xanthomonads strains). The different suspensions were mixed with 250 mg of healthy pseudostem of banana (*Musa acuminata*) that was previously crushed in NaOH 0.5 M + 2% PVP buffer (plant tissue homogenized in suspension at a 1:10 w/v ratio), and the whole mixture was extracted as described in the section “Simplified DNA extraction protocols” and tested with the different LAMP assays.

### Inoculations in controlled conditions and sampling

Inoculations of the different bacteria on four-leaf stage Cavendish banana plants were performed in a climate chamber within a biosafety level 3 containment facility, with a 12-hour photoperiod, a thermoperiod of 28 °C day, 25 °C night, and 80% hygrometry. For BBD and BXW, one representative strain was inoculated; for Moko, six strains were inoculated for representing each specific Moko subgroup: IIB-3 (n=1), IIB-4 (n=2), IIA-6 (n=2), IIA-24 (n=1). Bananas were thus challenged by RUN62 (BBD), NCPPB4379 (BXW), RUN74 (Moko IIB-3), RUN586 (Moko IIB-4), RUN579 (Moko IIB-4), RUN20 (Moko IIA-24), RUN9 (Moko IIA-6) and RUN395 (Moko IIA-6). The inoculations were performed by infiltrating 1 ml of a bacterial suspension at a concentration of 1 x 10^8^ CFU/ml into the base of the pseudostem of the banana plants as described in [15]. Three plants were inoculated per strain, and symptom monitoring was conducted periodically over more than two months using a 0 to 4 rating scale [39]. Six plants received Tris buffer instead of bacterial suspension (controls). Two pieces of pseudostem - one just above the corm (lower part) and one above the lower piece (upper part), along with a corm sample (100 mg) for some plants, were collected from each inoculated plant, and DNA was extracted using both the NaOH-based simplified DNA extraction protocol and the DNeasy plant extraction Kit. LAMP assays were performed on all DNA extracts. Additionally, 100 mg of adjacent pieces of pseudostem and corm were sampled and homogenized in 10 ml of Tris buffer. Fifty microliters of the homogenate were plated in duplicates on dedicated semi-specific media (see “Bacterial strains and DNA extraction” section) to estimate target concentrations.

### Field surveys

The different LAMP assays were tested on naturally infected samples during field surveys organized in collaboration with national and transnational partners: International Institute of Tropical Agriculture (IITA), National Research and Innovation Agency (BRIN), Banana Board and Escuela Superior Politécnica del Litoral (ESPOL), in Uganda, Indonesia (Java), Jamaica and Ecuador, respectively (S1 Fig.). Samples were collected from symptomatic plants and healthy plants (controls). From each plant, a tissue piece was collected for the LAMP assay and an adjacent sample for a reference PCR assay or qPCR assay.

In Uganda, Xvm was surveyed in 17 different plots distributed among three locations: Hoima, Kayunga and Mukono (S1 Fig., A). A total of 46 stem samples, including a set of 12 healthy pseudostem samples collected as negative controls were collected and tested by LAMP and PCR [31] after the simplified NaOH-based DNA extraction methods.

In West Java, Indonesia, surveys were conducted in the Cianjur regency, in four different plots within the Cikalongkulon and Mande districts (S1 Fig., B). As *Ralstonia syzygii* subspecies *celebesensis* (BBD disease) and a devastating fungal pathogen (*Fusarium oxysporum* f. sp. *cubense* TR4 = Foc TR4) were both described in these areas, plants displaying symptoms potentially caused by these pathogens were investigated, as well as healthy plants. In addition, atypical symptoms, as well as those potentially induced by the banana stem borer, were also investigated. Thirty-two plants were sampled and a total of 71 samples, collected from fruit, central core, pseudostem, corm or peduncle, were analyzed by LAMP. Of these, 61 were also analyzed by qPCR (27), following the simplified NaOH-based DNA extraction method.

In Ecuador, surveys were conducted by ESPOL in three areas, Quevedo (Los Rios Province), Valencia (Los Rios Province) and El Carmen (Manabí Province) (S1 Fig., C). Twenty-three samples were collected from the pseudostem tissues of symptomatic banana and plantain plants, and DNA extracted using the simplified NaOH-based protocol, before being sent to the CIRAD UMR PVBMT laboratory for duplex MOKO-LAMP testing. In parallel, ESPOL performed a multiplex PCR [25,26] on DNA extracted using the ISB mDNA extraction method [40]. Additionally, some bacteria were isolated in Ecuador from infected samples on SMSA medium [41]. DNA was extracted from pure culture according to the method described in [42] and tested using both LAMP and the MOKO-specific multiplex PCR [26].

In Jamaica, eradication campaigns for Moko disease have been previously conducted and only three stem and corm samples from the region of Saint James were available for analysis using the duplex MOKO-LAMP assay (S1 Fig., D) after the NaOH-based DNA extraction method. DNA extraction was also performed on adjacent tissues pieces using the Qiagen DNeasy plant kit, and tested by both duplex-PCR [27] and multiplex PCR [25,26]. Ten pseudostem samples collected from a healthy banana plant located on the UWI Mona Campus in Jamaica were similarly analyzed.

### Statistics

The Wilcoxon signed-rank test was used to compare the effect of extraction methods (NaOH-based simplified DNA extraction protocol versus DNeasy plant extraction Kit) on LAMP responses (TTR) using the R statistical software (version 4.1.1 [2021-08-10]; R Development Core Team,4, Vienna, Austria).

## Results

### Selection of bacterial targets and LAMP assays design

Seventy-five candidate genes were found for BBD. After a screening based on DNA fragment size selection (elimination of fragment < 300 bp), the elimination of phage related fragments and the specificity results on the different NCBI databases, two candidate targets were retained, identified within the genome of the strain ICMP10001 (R229): BBDv6_60001 (1245 bp) and BBDv6_50108 (1665 bp) encoding for a protein of unknown function and a putative hemolysin activation/secretion protein, respectively (S5 Table). For BXW, 50 candidate sequences were found. From the same screening process, two sequences were retained from the Xvm genome NCPPB4379: KWO_005740 (405 bp) and KWO_019835 (369 pb), encoding for a histone-like nucleoid-structuring protein and a hypothetical protein, respectively. For MOKO IIA, two genes within the genome of the IIA-24 strain B50 were selected among 28 candidate genes: RALB5v2_2960030 (639 bp) and RALB5v2_200021 (942 bp), encoding a conserved protein of unknown function and a plasmid replication region DNA-binding protein, respectively. Interestingly, RALB5v2_2960030 was also present in all IIB-3 genomes. For MOKO IIB, no genes were found to be common to both IIB-3 and IIB-4 strains while being absent in non-target genomes. However, 35 candidate genes were shared specifically by all IIB-4 strains while absent in the IIB4-NPB strains. From the screening process, RSPO_c00972 (540 bp) and RSPO_c00967 (645 bp), encoding for a conserved zinc ribbon domain-containing protein and a conserved protein of unknown function, respectively, were selected from the genome of the IIB-4 strain PO82.

One LAMP primer set was successfully designed for KWO_005740 and RALB5v2_2960030, two LAMP primer sets for BBDv6_60001, KWO_019835, RALB5v2_200021, RSPO_c00972, three LAMP primer sets for BBDv6_50108 and four primer sets for RSPO_c00967. LAMP assays were run on a few sets of target strains and a “no Template Control” (NTC): RUN 62, RUN1314 and RUN1350 for BBD; NCPPB4387, XCM213B and BCC250 for BXW; RUN25, RUN49 and RUN96 for Moko. The LAMP primer set that gave the earliest TTR values and no signal for the NTC samples (no primer dimers) was selected. The final selected targets were BBDv6_50108, KWO_005740, RALB5v2_2960030 and RSPO_c00972, for BBD, BXW, MOKO II-A and MOKO II-B, respectively. The different LAMP primer sets selected for each pathogen are listed in Table 1. Interestingly, a stem primer, manually designed between the B1 and F1 regions targeted by the LAMP assay as described by [43] successfully speeded up the reaction when added to the BXW LAMP primer set.

**Table 1.**
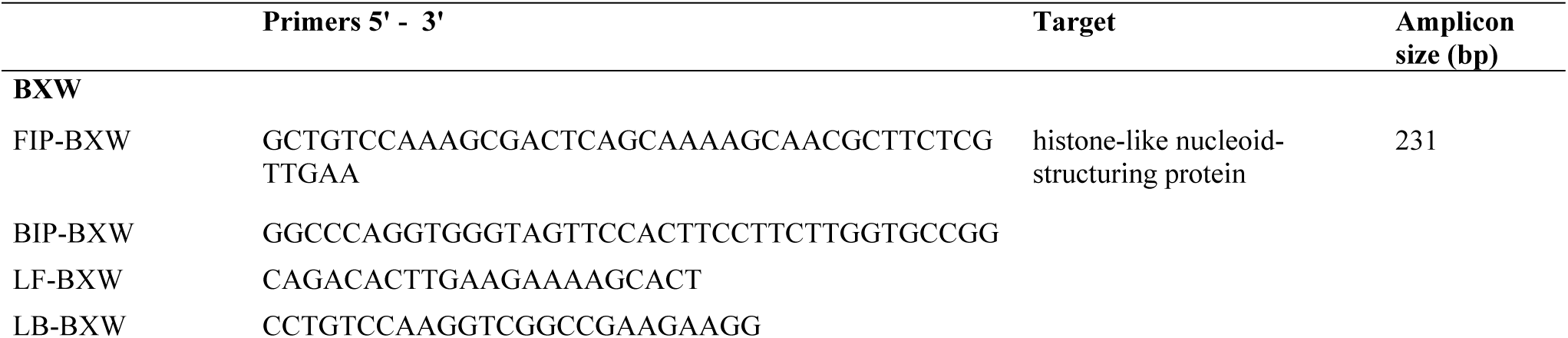
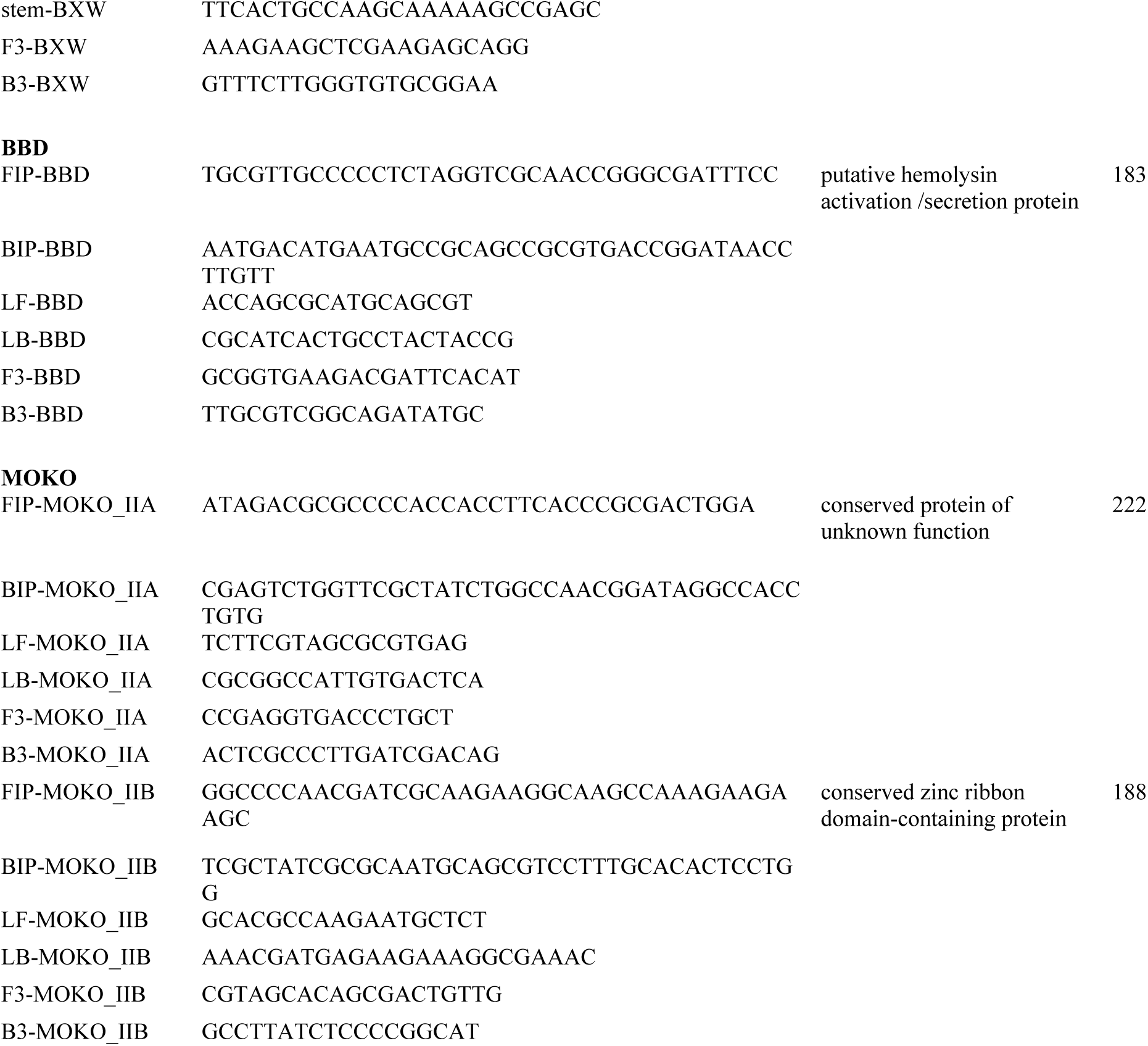
LAMP Primers designed in this study.

### Optimization and profiles of the LAMP assays

From the gradient tests, the best TTR values for BBD-LAMP assay were obtained for the range 65-67 °C, and the standard temperature 65 °C was selected (Fig 1, A). For the MOKO-LAMP assay, the best values were obtained for 66-68 °C, and the temperature of 67 °C was selected (Fig 1, B). For the BXW-LAMP assay, the first assays made with the standard primer concentration (1X) showed late positive signals compared to the other LAMP systems. Doubling the concentration allowed for earlier positive signal. The best results were obtained for 66 and 67 °C (Fig 1, C). A LAMP reaction temperature of 66 °C was then chosen.

**Fig. 1.**
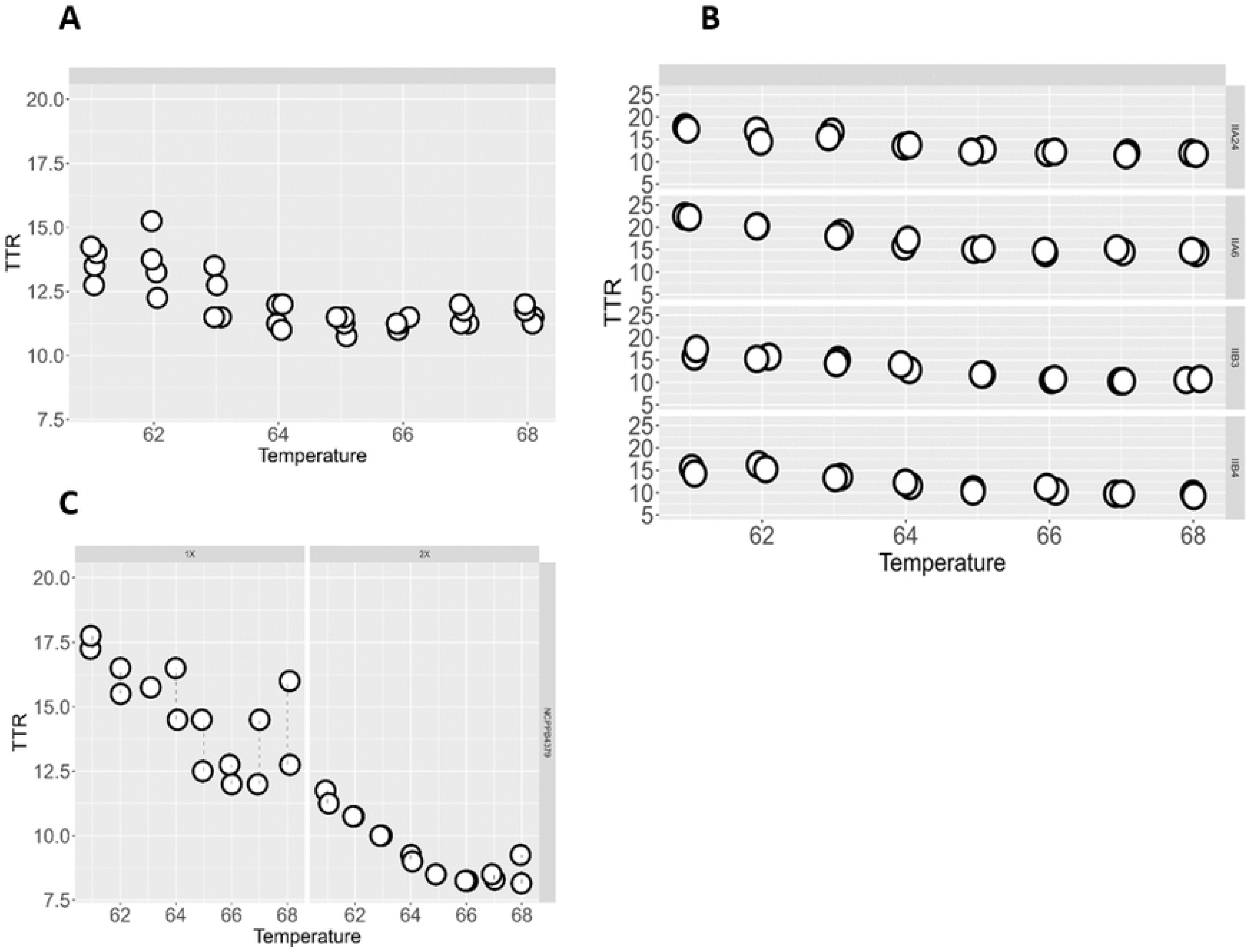
TTR signals according to different LAMP reaction temperatures (gradient from 62 °C to 68 °C). (A) BBD-LAMP, (B) MOKO-duplex LAMP on four strains representative of the different sequevars, (C) BXW-LAMP at two primer concentrations: 1X and 2X. The TTR values are the “min” values i.e. the values corresponding to the take-off of the fluorescence.

The different LAMP assays profiles are illustrated in Fig 2. A sample was considered positive if TTR<30 and Annealing Temperature (Ta) peaks reached a minimal fluorescence threshold value of 1500 RFU (Relative Fluorescence Unit), and displayed a specific Ta peak: 89.5 °C <Ta< 91 °C for BXW, 90 °C <Ta< 92 °C for BBD and at least one peak between 94 °C-96 °C (IIA marker amplification) or 90 °C-91.5 °C (IIB marker amplification) for Moko. The IIA-24 strains were expected to display both melting peaks. However, the IIB peak tended to take over the IIA peak. The latter may be reduced or even absent. We tried unsuccessfully to optimize the assay by adjusting the ratio of the two primer sets (data not shown). Nevertheless, the presence of at least one peak was sufficient to diagnose Moko disease. A simplified and automatic output of results was programmed on the Genie II (Fig 2).

**Fig. 2.**
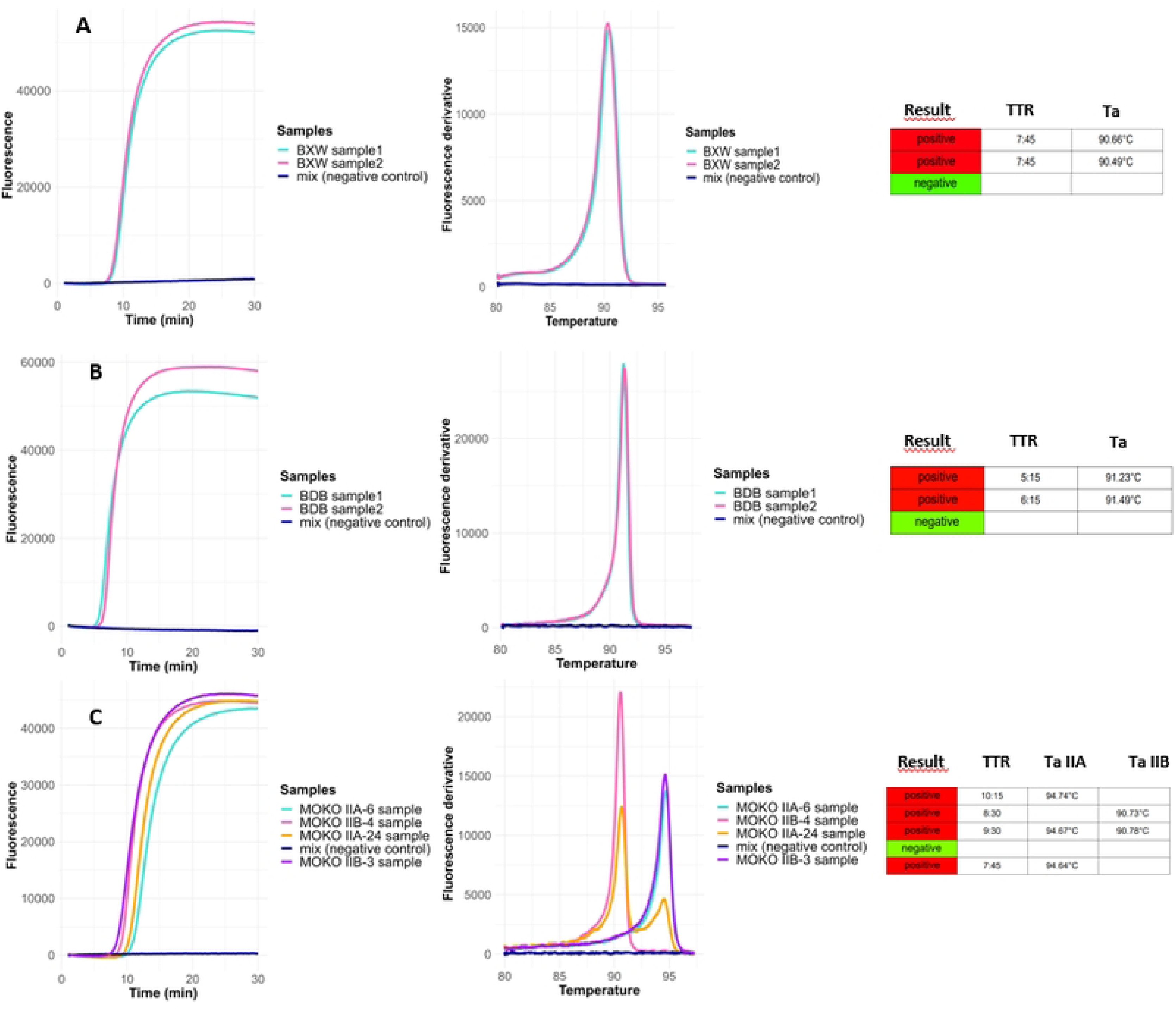
Profiles of the different LAMP assays on Genie II instrument. Left: Amplification curves, center: Temperature annealing peaks, right: automatic readouts. (A) BXW-LAMP assay, (B) BBD-LAMP assay, (C) MOKO duplex LAMP assay.

### Specificity

#### In silico analysis

The specificity of each LAMP primer set was verified using a wide collection of target and non-target genomes. For the BXW-LAMP assay, the target region was detected in all Xvm genomes (n = 24) present in the database with 100% identity to query sequence, corresponding to one variant (S6 Table). No matches with the query sequence were observed in most of the non-target genomes (n = 7,381), including 86 genomes of *Xanthomonas vasicola* pathovars. However, some others non-target Xanthomonads genomes (n = 78) showed low identity level, ranging from 84.7% to 73.3% (variant 2 to 12), which would likely prevent the LAMP reaction from working.

For the BBD-LAMP assay, the target region matched the eight genomes of *R. syzygii* subsp. *celebesensis* available in the database with 100% identity. No matches were obtained for the 9,960 non-target genomes, including closely related subspecies of *R. syzygii* (S7 Table).

The IIA target of the duplex MOKO-LAMP assay was detected in all the Moko strains belonging to the sequevars IIA-6, IIA-24 and IIB-3 present in the database (n = 27) with 100 % identity (one variant), excepted in four strains belonging to IIB-3: PD1446, UW75, UW136 and RUN 579 (S8 Table). These strains were all isolated in the 1950s in Costa Rica from *Heliconia sp*. except UW75 strain that was isolated from *M. acuminata*. No matches were obtained with the 9,895 non-target genomes.

The IIB target DNA was detected in all IIB-4 (n=42) and IIA-24 (n = 6) strains with 100 and 98.7% identity, respectively (two variants). Most of the non-target strains (n = 9,904), including the IIB-4 NPB genomes (n=28), displayed no significant match with the IIB-4 target DNA. It should be noted that the non-pathogenicity to banana of most of the IIB-4 NPB strains has been verified [15,44]. Moreover, all these genomes contained the amplicon described by Cellier et al [27], specific of the IIB-4 NPB strains. The IIB target DNA was absent in two strains UA-1579 and UA-1609, even though they were considered as Moko-inducing strains [45,46]. Meanwhile, we detected in both genomes the IIB-4NPB amplicon [27]. Fourteen genomes, mostly corresponding to other *Ralstonia* species (*R. pickettii*, *R. thomasii*), matched the IIB target DNA, with identity ranging from 94.3% to 94.9% (two variants). The accumulation of mutations could compromise LAMP efficiency, notably one mutation at the 3’ extremity of F3 and two closed mutations in FIP-F2 or FIP-F1c [47].

#### Analytical specificity

The experimental data confirmed the *in silico* results. One hundred percent inclusivity was obtained for the BBD and BXW-LAMP assays tested on the collection of respective target strains (Table 2 and Table S1 and S3 Tables). For Moko disease, all 41 target strains tested positive, except RUN579, the only strain available in our collection among the four strains isolated in the 1950s in Costa Rica (Table 2 and S2 Table). However, no symptoms were observed after inoculating this strain on Cavendish banana plants in our conditions.

**Table 2.**
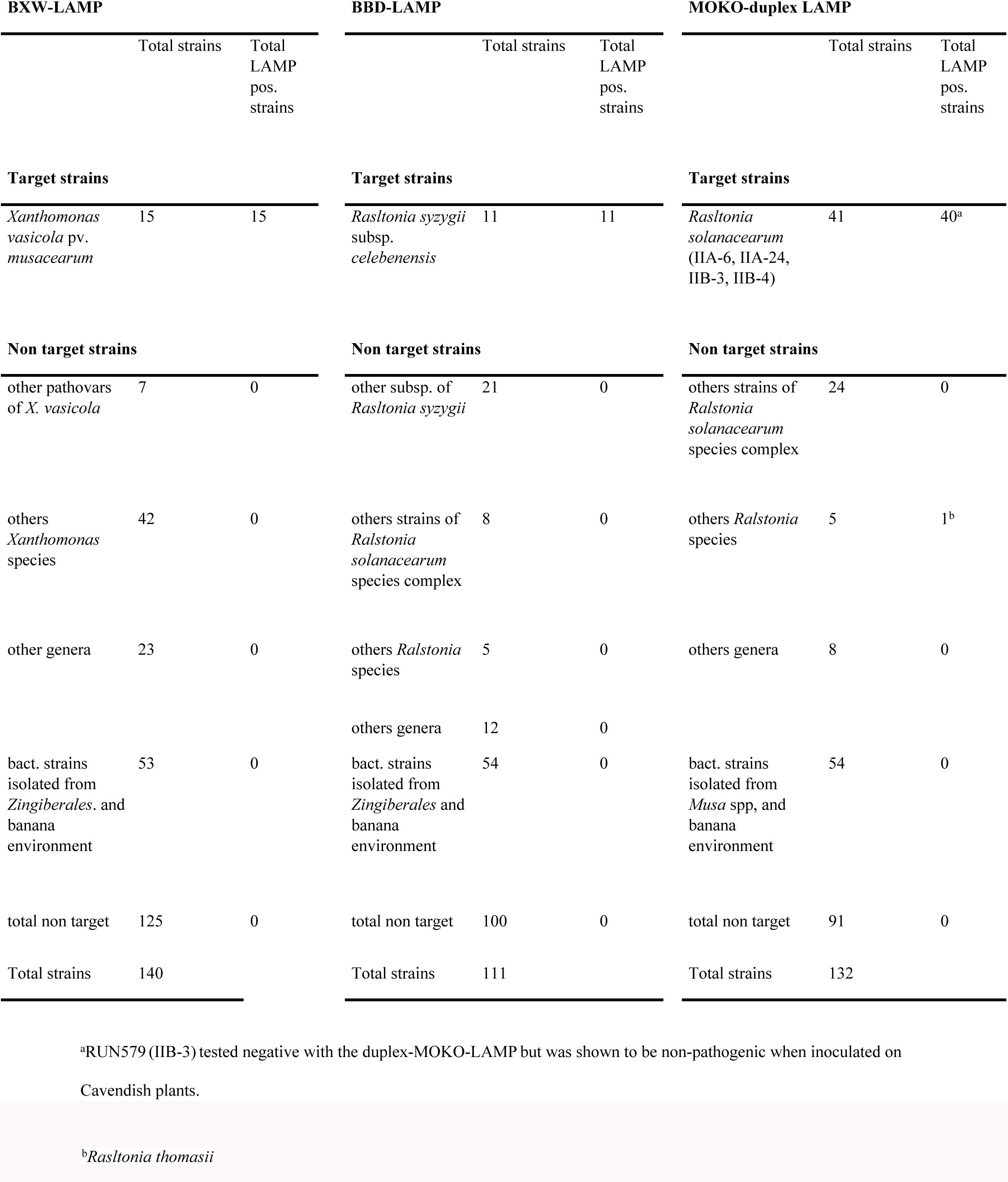
Specificity of the LAMP assays evaluated on collection of target and non-target strains.

All non-target strains (n = 125) tested negative with the BXW-LAMP assay, including closely related strains such as other pathovars of *X. vasicola* (e.g., pv. *holcicola*, pv. *vasculorum*), the closely related species *Xanthomonas oryzae* pv. *oryzae*, as well as strains isolated from Zingiberales and the banana environment (soil/rhizosphere) (n = 53). Similarly, no LAMP signal was obtained with the BBD-LAMP assay when tested on the non-target strain collection (n = 100), which included the other subspecies of *R. syzygii*, *indonesiensis* and *syzygii*, as well as strains isolated from Zingiberales and the banana environment. For Moko, only a single strain, belonging to *R. thomasii,* produced a delayed amplification signal out of the 91 non-target strains tested. These included various sequevars from subgroups IIA and IIB (1, 2, 7, 28, 35, 41, 4NPB), along with 54 strains isolated from *Musa* spp. and the banana environment (Table 2; S1–S3 Tables).

### Analytical sensitivity

The limit of detection (LOD) (100 % positive responses) was first evaluated on serial dilutions of DNA(0.1 to 0.00001 ng/µl). All LOD values, except for the MOKO IIA-24 strain RUN24, were 1 pg/µl, which corresponds to estimated values of genome copies ranging from 165 to 175 genome copies/µl according to the size of the genomes (Table 3). For RUN24, the LOD was ten-fold lower corresponding to 16 copies/µl. When performed on spiked banana tissues (Fig 3), a LOD values of 10^4^ CFU/ml was observed for the BBD and BXW-LAMP assays and the IIA-24 MOKO strain, and 10^5^ CFU/ml for the other three MOKO strains.

**Fig. 3.**
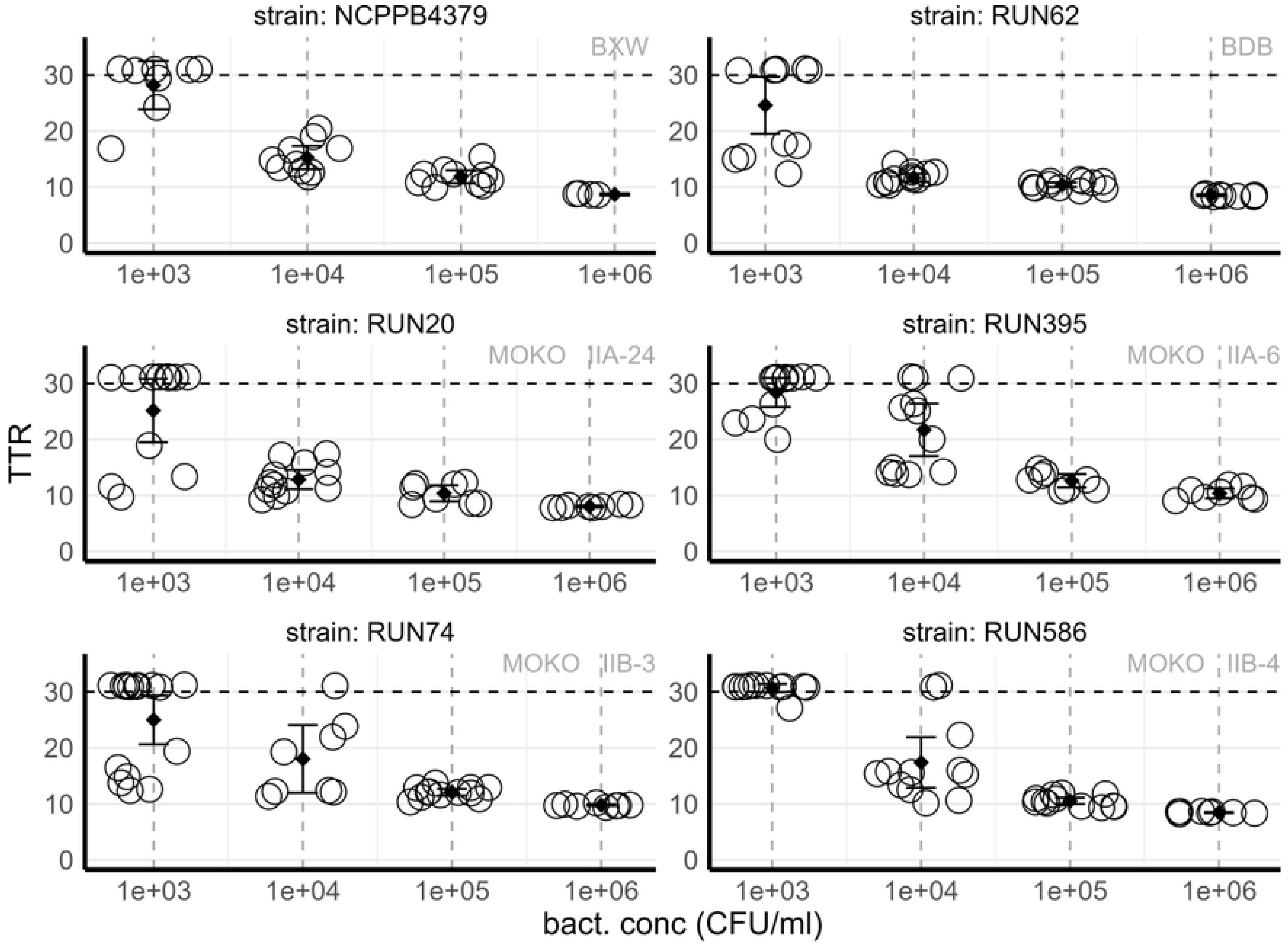
Sensitivity of the LAMP assays on heathy tissues spiked with different concentrations of bacterial suspensions (from 10^3^ to 10^6^ CFU/ml). DNA was extracted with the simplified NaOH-based DNA extraction method and tested with the appropriate LAMP assay (8-12 replicates according to the dilutions).

**Table 3.**
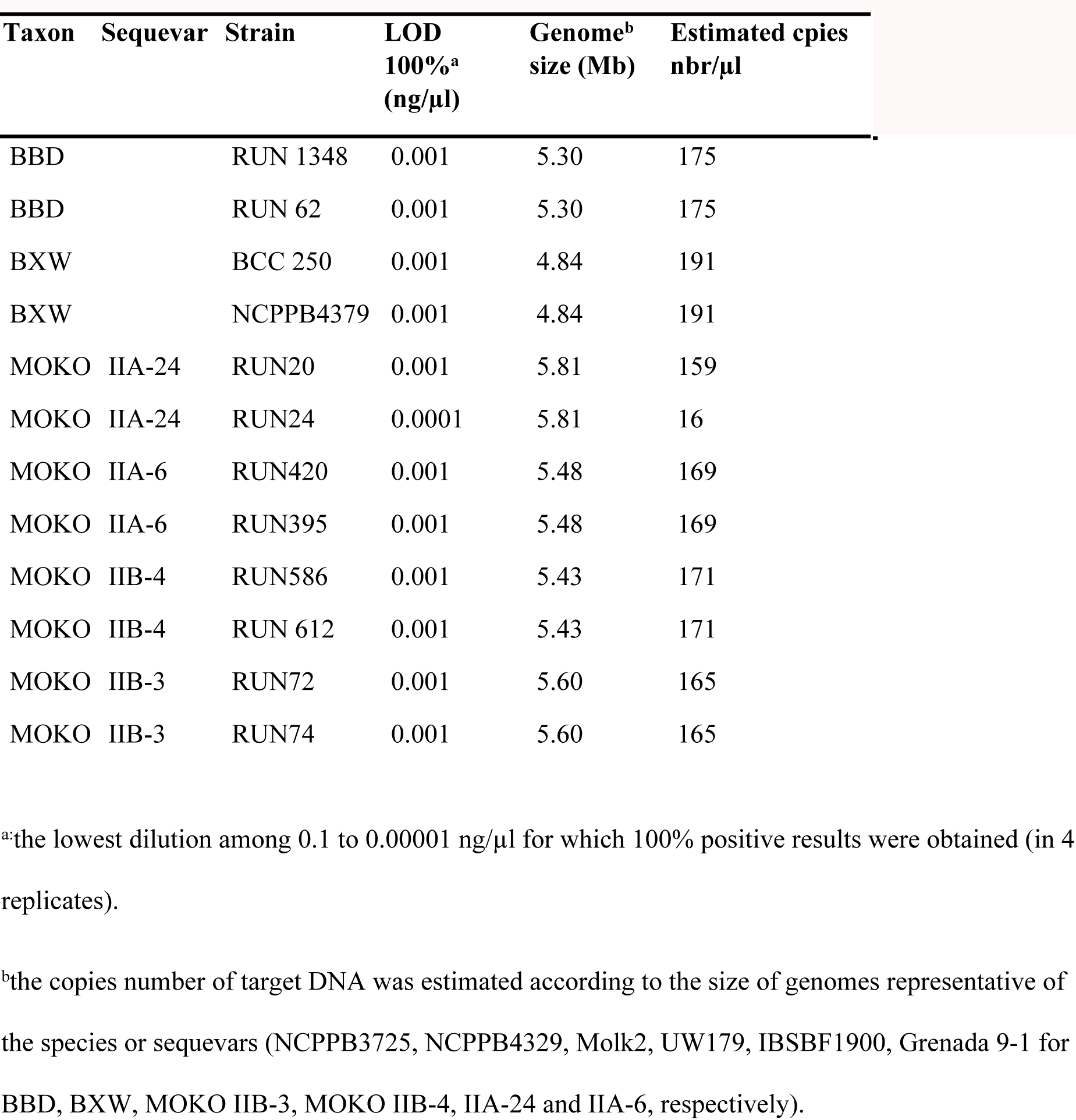
Limit of detection (LOD) obtained for various BXW, BBD and MOKO strains evaluated on serially diluted DNA.

### LAMP detection from greenhouse-inoculated plants

All inoculated strains, except the Moko strains RUN9 and RUN579, caused typical vascular symptoms (yellowing, necrosis and wilting) in the test plants and reached the maximum severity rating of four for at least one inoculated plant Fig 4). No symptoms were observed on the plants inoculated with the Moko strain IIB-4 RUN579. The Moko strain RUN9 caused only mild symptoms on the inoculated plants with reduced size compared to the uninoculated controls, slight leaf wilting, and yellowing and necrosis of oldest leaves. Moreover, internal symptoms such as necrosis on transverse sections of the roots and corm were associated with this “mild symptom profile”. Additionally, the Moko strain RUN20 caused variable symptoms depending on the plants: typical, mild (similar to those induced by RUN9), or none. No symptoms were observed on the control plants.

**Fig. 4.**
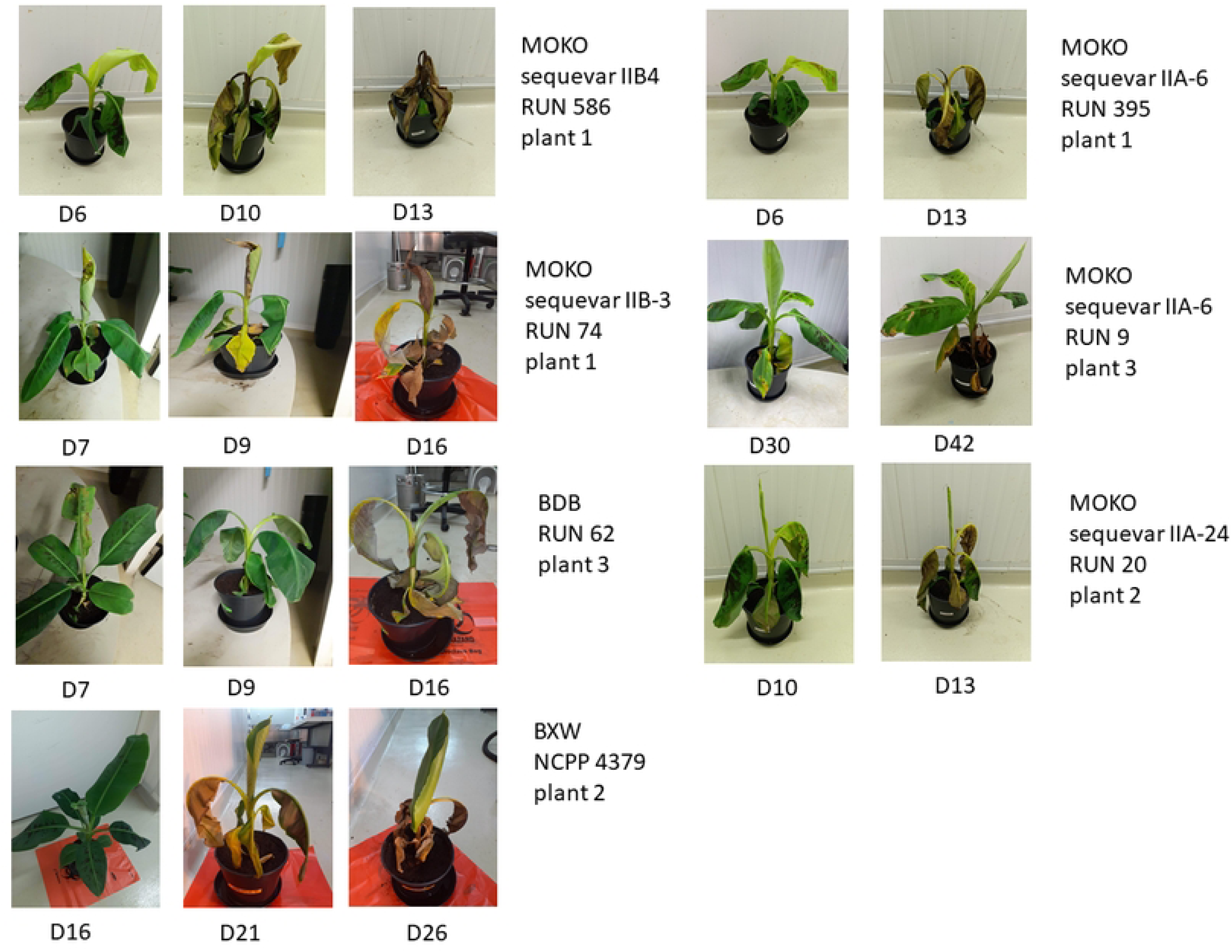
Symptoms observed over time after inoculation of strains belonging to the species *R. solanacearum* (Moko disease), *R. syzygii* subsp. *celebesensis* (BBD), and *X. vasicola* pv*. musacearum* (BXW). “D” followed by a number refers to the day after inoculation when the observation occurred.

Differences were observed among the strains regarding the timing of symptoms onset and the number of infected plants. All plants inoculated with BBD, Moko IIB-3 and IIB-4 strains exhibited symptoms within 15 days whereas *Xanthomonas* wilt progressed more slowly, with 100% of the plants displaying symptoms after 30 days. Among the Moko IIA strains, RUN595 induced typical symptoms on all plants, but with delayed onset compared to the IIB strains. Plants inoculated with RUN20 and RUN9 that developed mild symptoms showed a delayed onset of symptom expression. (S2 Fig.).

No LAMP signal was detected, and no bacteria were recovered from the control plants. Similarly, no duplex-MOKO LAMP signal was detected in any plants inoculated with the IIB-4 strain RUN579 and accordingly, no typical bacterial colonies were recovered from their tissues (data not shown).

Conversely, for the other strains, each LAMP assay successfully amplified the target bacteria directly from pieces of symptomatic tissue sampled in the pseusdostem (lower and upper part) of the infected plants, or in the corm sampled from some plants, after either simplified DNA extraction or kit-based extraction (Fig 5). No significant differences were obtained between the two extraction methods (p= 0.3738). High bacterial concentrations (≈ 10⁷–10⁸ CFU/ml) and early TTR values (≤ 10) were obtained for all plants displaying typical symptoms, regardless of the banana disease. The identity of bacterial colonies was confirmed through specific LAMP assays. Bacteria were isolated from all three plants inoculated with the Moko strain RUN9, displaying mild symptoms, with bacterial concentrations ranging between 1.9 × 10^5^ – 1.7 × 10^7^ CFU/ml, depending on the plant and tissues sampled. LAMP detection was achieved even in the plant n° 2, for which the concentration of isolated bacteria ranged between 4 × 10^5^-8.9 × 10^5^ CFU/ml.

**Fig. 5.**
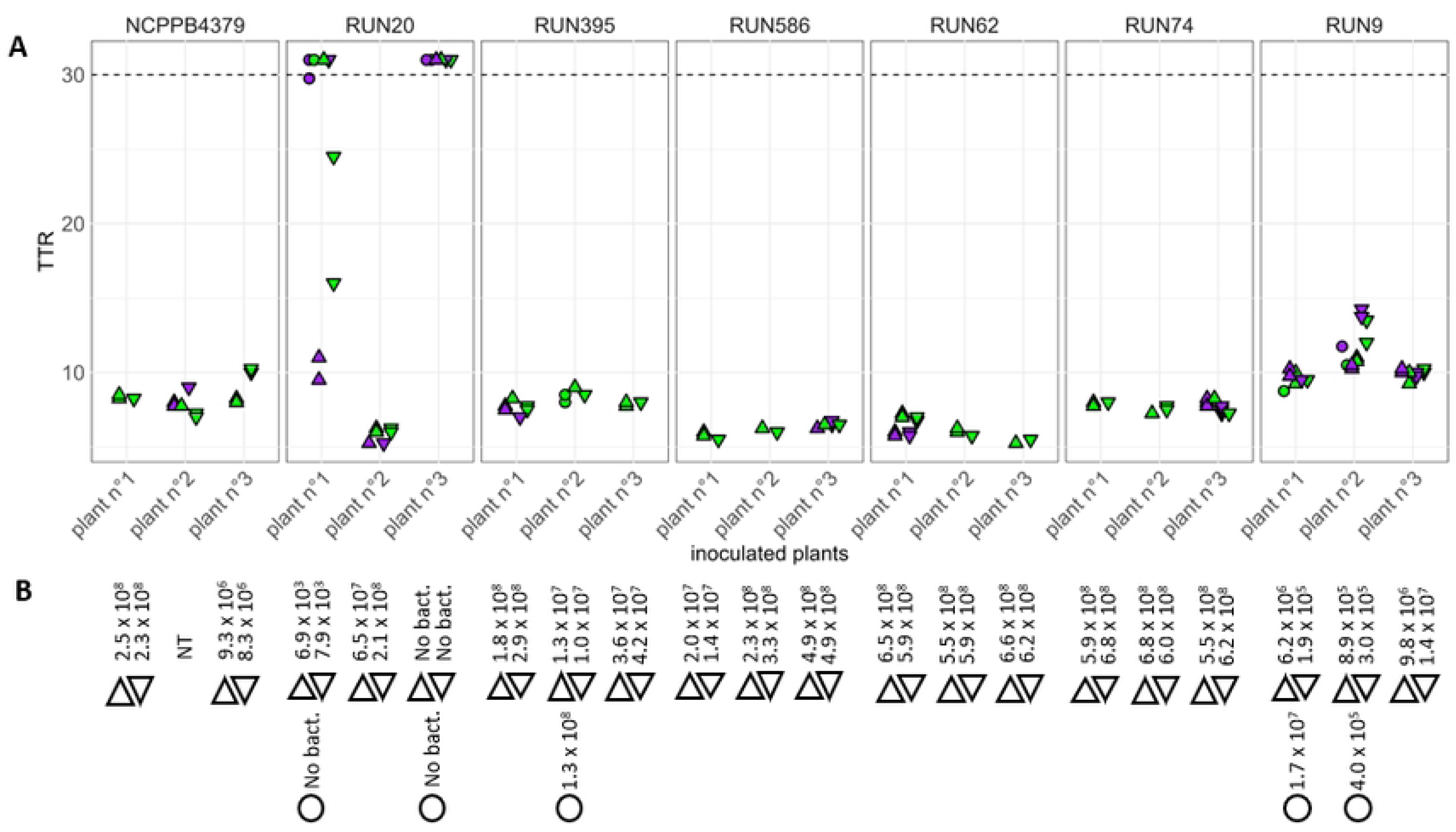
LAMP signals (A) and bacterial concentrations (B) obtained from inoculated banana plants. ∆: pseudostem, upper part, ∇: pseudostem lower part, ○: corm. For LAMP tests, the plant tissues were extracted with an extraction kit 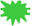 or the simplified NaoH-based extraction 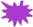.

For Moko IIA-24 strain RUN20, no LAMP signal was observed and no bacterial colonies could be isolated from the asymptomatic plant n°3. Additionally, only a few TTR signals were observed for the plant n°1 displaying a “mild symptom” profile, and correspondingly, the bacterial concentration was low (6.9–7.9 × 10³ CFU/ml).

### LAMP detection from naturally infected field samples

The different LAMP assays were tested on naturally infected field samples from different countries, representing contrasted disease contexts, to evaluate their detection efficiency on symptomatic banana plants, in comparison with reference PCR and qPCR assays.

#### Xanthomonas wilt

Thirty-two samples mostly collected from the pseudostem tissues of banana plants, displaying typical symptoms of BXW, tested positive with the BXW-LAMP assay (Table 4). Bacterial ooze was sometimes observed in pseudostem transverse sections. Various Ugandan banana cultivars investigated included Sukali Ndiizi, Gros Michel, Kayinja (Pisang Awak group), Nakabululu, plantain, and different cooking banana types. The positive samples were distributed across all plots visited in the three localities, except plot 2. In the plot 2, the BXW-LAMP negative samples were taken from two declining banana plants displaying internal symptoms characteristic of *Fusarium* wilt. Additionally, 12 healthy samples tested all negative using the BXW-LAMP assay. The TTR ranged between 5.75 and 13.5 min (median: 7.87 min), and the Ta between 90.74 and 90.94 °C. No signal was obtained for the NTC controls. LAMP assays gave results highly consistent with the reference PCR assay [31], performed on the same extracts (99% concordance, with only one sample tested negative with PCR while being LAMP-positive).

**Table 4.**
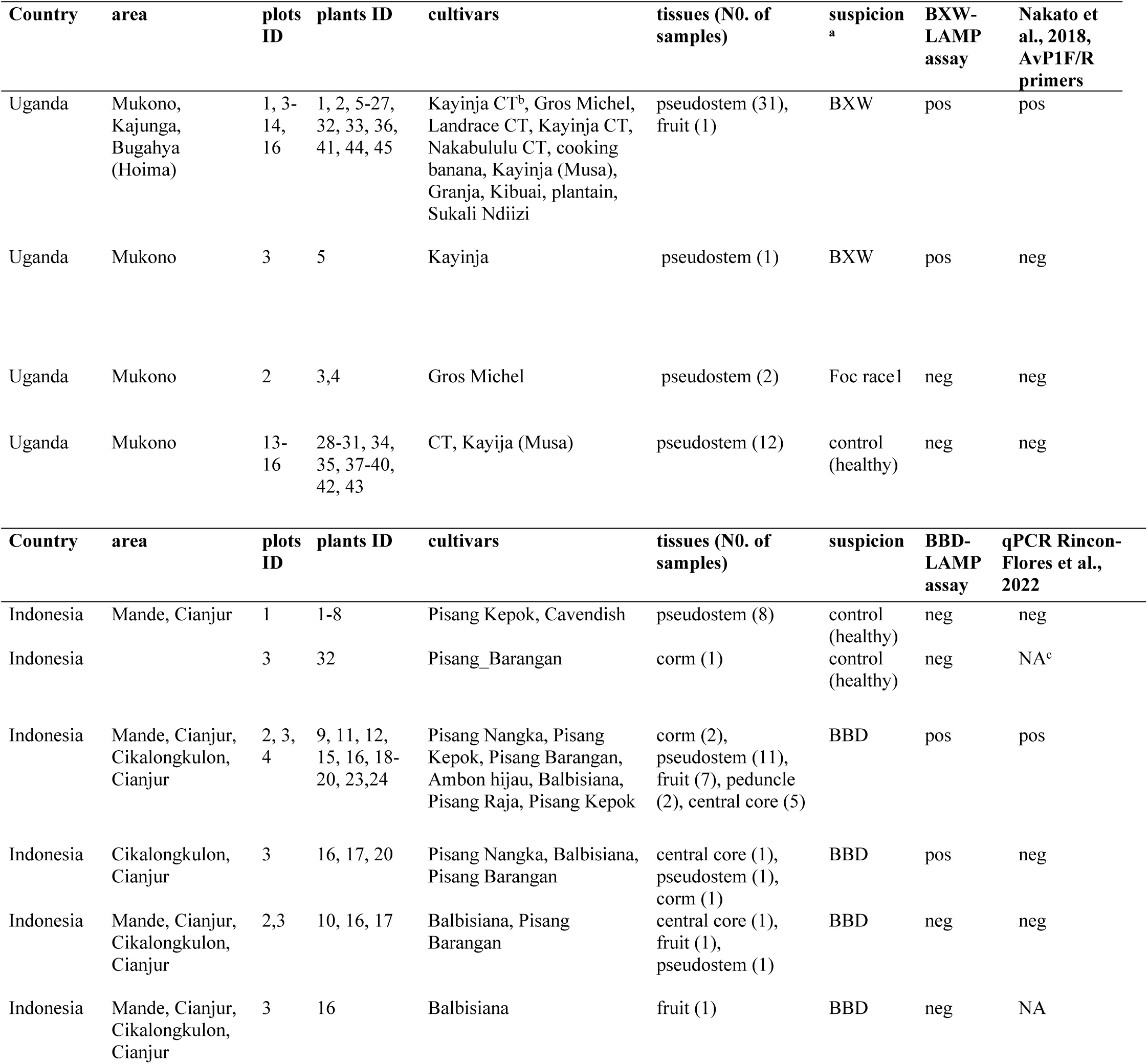
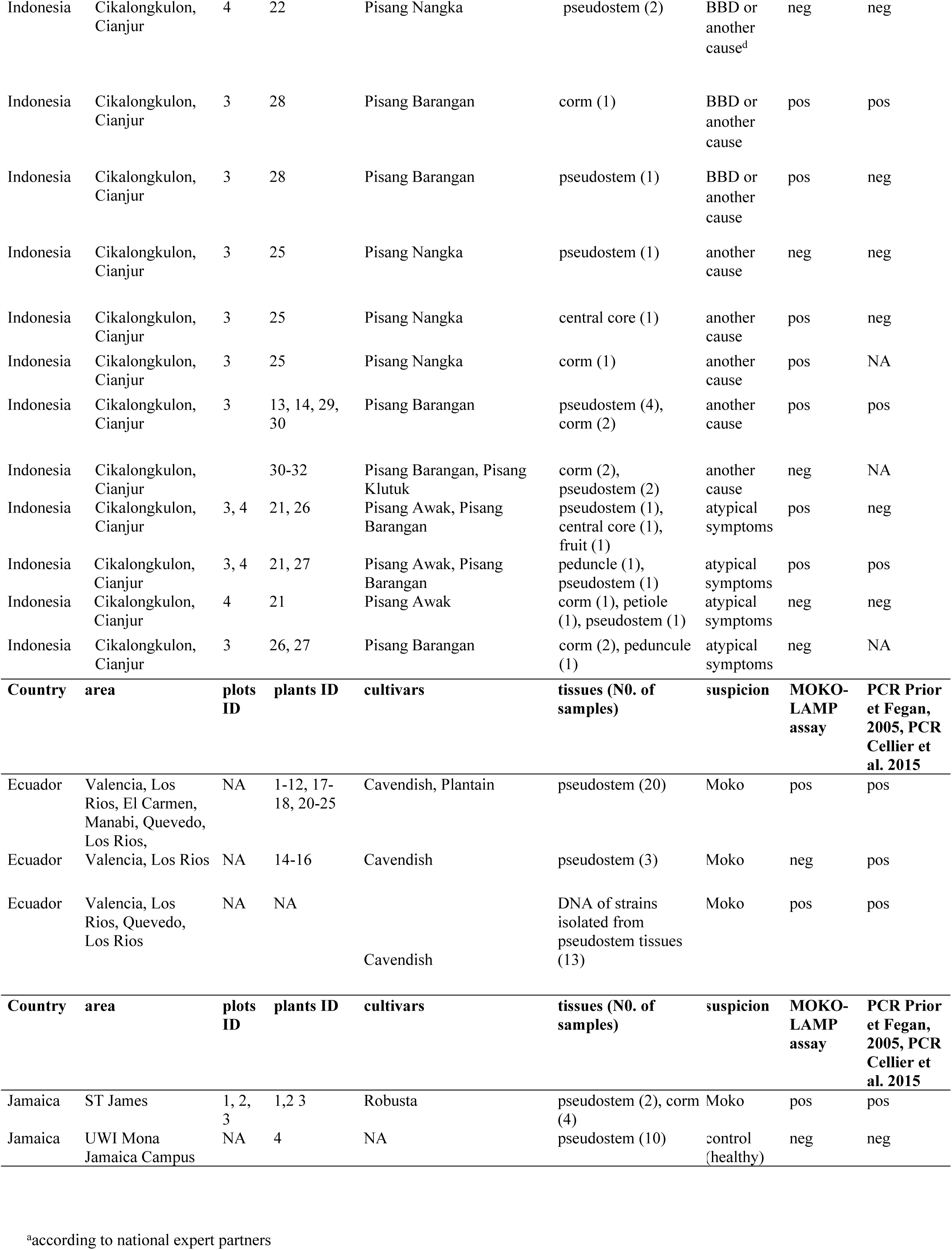
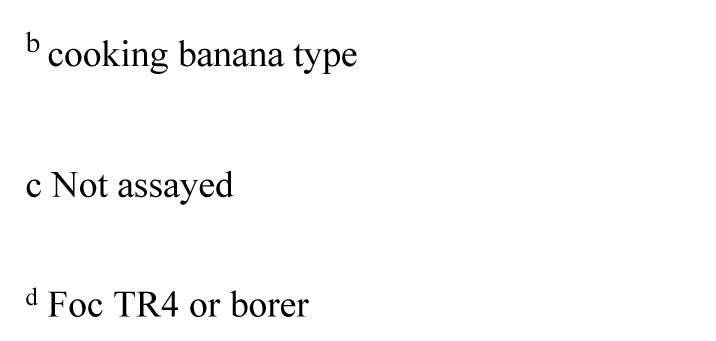
Comparison of LAMP and PCR/qPCR conducted on field samples.

| Country | area | plots ID | plants ID | cultivars | tissues (N0. of samples) | suspicion<sup>a</sup> | BXW-LAMP assay | Nakato et al., 2018, AvP1F/R primers |
| --- | --- | --- | --- | --- | --- | --- | --- | --- |
| Uganda | Mukono, Kajunga, Bugahya (Hoima) | 1, 3-14, 16 | 1, 2, 5-27, 32, 33, 36, 41, 44, 45 | Kayinja CT<sup>b</sup>, Gros Michel, Landrace CT, Kayinja CT, Nakabululu CT, cooking banana, Kayinja (Musa), Granja, Kibuai, plantain, Sukali Ndiizi | pseudostem (31), fruit (1) | BXW | pos | pos |
| Uganda | Mukono | 3 | 5 | Kayinja | pseudostem (1) | BXW | pos | neg |
| Uganda | Mukono | 2 | 3,4 | Gros Michel | pseudostem (2) | Foc race1 | neg | neg |
| Uganda | Mukono | 13-16 | 28-31, 34, 35, 37-40, 42, 43 | CT, Kayija (Musa) | pseudostem (12) | control (healthy) | neg | neg |
| Country | area | plots ID | plants ID | cultivars | tissues (N0. of samples) | suspicion | BBD-LAMP assay | qPCR Rincon-Flores et al., 2022 |
| Indonesia | Mande, Cianjur | 1 | 1-8 | Pisang Kepok, Cavendish | pseudostem (8) | control (healthy) | neg | neg |
| Indonesia |  | 3 | 32 | Pisang_Barangan | corm (1) | control (healthy) | neg | NA<sup>c</sup> |
| Indonesia | Mande, Cianjur, Cikalongkulon, Cianjur | 2, 3, 4 | 9, 11, 12, 15, 16, 18-20, 23,24 | Pisang Nangka, Pisang Kepok, Pisang Barangan, Ambon hijau, Balbisiana, Pisang Raja, Pisang Kepok | corm (2), pseudostem (11), fruit (7), peduncle (2), central core (5) | BBD | pos | pos |
| Indonesia | Cikalongkulon, Cianjur | 3 | 16, 17, 20 | Pisang Nangka, Balbisiana, Pisang Barangan | central core (1), pseudostem (1), corm (1) | BBD | pos | neg |
| Indonesia | Mande, Cianjur, Cikalongkulon, Cianjur | 2,3 | 10, 16, 17 | Balbisiana, Pisang Barangan | central core (1), fruit (1), pseudostem (1) | BBD | neg | neg |
| Indonesia | Mande, Cianjur, Cikalongkulon, Cianjur | 3 | 16 | Balbisiana | fruit (1) | BBD | neg | NA |

| Indonesia | Cikalongkulon, Cianjur | 4 | 22 | Pisang Nangka | pseudostem (2) | BBD or another cause<sup>d</sup> | neg | neg |
| --- | --- | --- | --- | --- | --- | --- | --- | --- |
| Indonesia | Cikalongkulon, Cianjur | 3 | 28 | Pisang Barangan | corm (1) | BBD or another cause | pos | pos |
| Indonesia | Cikalongkulon, Cianjur | 3 | 28 | Pisang Barangan | pseudostem (1) | BBD or another cause | pos | neg |
| Indonesia | Cikalongkulon, Cianjur | 3 | 25 | Pisang Nangka | pseudostem (1) | another cause | neg | neg |
| Indonesia | Cikalongkulon, Cianjur | 3 | 25 | Pisang Nangka | central core (1) | another cause | pos | neg |
| Indonesia | Cikalongkulon, Cianjur | 3 | 25 | Pisang Nangka | corm (1) | another cause | pos | NA |
| Indonesia | Cikalongkulon, Cianjur | 3 | 13, 14, 29, 30 | Pisang Barangan | pseudostem (4), corm (2) | another cause | pos | pos |
| Indonesia | Cikalongkulon, Cianjur |  | 30-32 | Pisang Barangan, Pisang Klutuk | corm (2), pseudostem (2) | another cause | neg | NA |
| Indonesia | Cikalongkulon, Cianjur | 3, 4 | 21, 26 | Pisang Awak, Pisang Barangan | pseudostem (1), central core (1), fruit (1) | atypical symptoms | pos | neg |
| Indonesia | Cikalongkulon, Cianjur | 3, 4 | 21, 27 | Pisang Awak, Pisang Barangan | peduncle (1), pseudostem (1) | atypical symptoms | pos | pos |
| Indonesia | Cikalongkulon, Cianjur | 4 | 21 | Pisang Awak | corm (1), petiole (1), pseudostem (1) | atypical symptoms | neg | neg |
| Indonesia | Cikalongkulon, Cianjur | 3 | 26, 27 | Pisang Barangan | corm (2), peduncle (1) | atypical symptoms | neg | NA |
| Country | area | plots ID | plants ID | cultivars | tissues (N0. of samples) | suspicion | MOKO-LAMP assay | PCR Prior et Fegan, 2005, PCR Cellier et al. 2015 |
| Ecuador | Valencia, Los Rios, El Carmen, Manabi, Quevedo, Los Rios, | NA | 1-12, 17-18, 20-25 | Cavendish, Plantain | pseudostem (20) | Moko | pos | pos |
| Ecuador | Valencia, Los Rios | NA | 14-16 | Cavendish | pseudostem (3) | Moko | neg | pos |
| Ecuador | Valencia, Los Rios, Quevedo, Los Rios | NA | NA | Cavendish | DNA of strains isolated from pseudostem tissues (13) | Moko | pos | pos |
| Country | area | plots ID | plants ID | cultivars | tissues (N0. of samples) | suspicion | MOKO-LAMP assay | PCR Prior et Fegan, 2005, PCR Cellier et al. 2015 |
| Jamaica | ST James | 1, 2, 3 | 1,2 3 | Robusta | pseudostem (2), corm (4) | Moko | pos | pos |
| Jamaica | UWI Mona Jamaica Campus | NA | 4 | NA | pseudostem (10) | control (healthy) | neg | neg |

<sup>a</sup>according to national expert partners
<sup>b</sup> cooking banana type
<sup>c</sup> Not assayed
<sup>d</sup> Foc TR4 or borer

#### Banana Blood Disease

The BBD-LAMP assay was successfully tested in three out of the four investigated plots in Indonesia, in various plant organs including fruit, central core, pseudostem, corm, and peduncle, as well as in different banana varieties such as Pisang Nangka, Pisang Kepok, Pisang Barangan, Ambon Hijau, and *M. Balbisiana* (Table 4). The LAMP TTR values ranged from 5.5 to 29.5 min (median: 11.62 min), while Ta values were very stable (90.75 to 91.42 °C). The reference qPCR assay [30], performed on the same simplified extracts, gave clear though slightly delayed signals compared to LAMP, with Ct values ranging from 19.29 to 38.83 (median: 30.31). Additionally, eight samples that tested positive with the LAMP assay were negative with the qPCR assay, likely due to the presence of inhibitors in the samples. However, a quite high concordance rate of 87% was observed between LAMP and the reference qPCR. Interestingly, the BBD-LAMP assay successfully detected *R. syzygii* subsp. *celebesensis* in six banana plants that also tested positive for the Foc TR4 pathogen when using a TR4-specific LAMP assay, highlighting the assay’s selectivity in mixed infections (data not shown). Symptoms initially attributed to other causes such as weevil borers or Fusarium wilt, may still test positive for BBD, based on LAMP and qPCR results. Similarly, atypical symptoms may be caused either by BBD or by another biotic agent (Table 4).

#### Moko disease

The MOKO-LAMP assay worked successfully when tested in two disease contexts. In Ecuador, it detected MOKO IIB-4 in 20 pseudostem samples collected in three areas: Valencia (Los Ríos), Quevedo, (Los Ríos), and El Carmen (Manabí). The TTR values ranged from 7 to 21.25 min (median: 12.75 min), with Ta values ranging from 90.3 to 90.92 °C, a signature of the IIB-4 sequevar. The Musa multiplex PCR assay [25,26], was also performed in Ecuador on the same samples, and all samples tested positive, showing a concordance of 87.5% with the LAMP results (three samples among 23 were LAMP-negative while PCR-positive). All 13 DNA extracted from strains isolated from pseudostem tissues tested positive with both molecular assays. The PCR profile obtained with the Musa multiplex assay confirmed the exclusive presence of the *R. solanacearum* sequevar IIB-4 in Ecuador.

In Jamaica, MOKO-LAMP successfully amplified the IIA marker in all samples from three plots in Saint James, like the reference PCR assay [27] and the Musa multiplex PCR assay [25,26] did, while healthy samples tested negative. LAMP and reference PCR were thus fully concordant. The PCR profile obtained with the Musa multiplex PCR assay corresponded to that of sequevar IIA-6.

## Discussion

A precise and efficient diagnostic method is essential for effective disease control and outbreak monitoring. Diagnostics are key to early detection, allowing for timely intervention and management, which can greatly limit pathogen spread and reduce disease impact. In this study we developed and validated reliable LAMP assays for the diagnosis of three major bacterial vascular diseases of banana, Moko Disease, Blood Disease of Banana (BBD) and Banana *Xanthomonas* Wilt (BXW). These tools will be useful for direct detection of the responsible pathogens from banana tissues in the field or outside a traditional laboratory environment.

Access to complete genomic data has advanced the use of comparative genomics to identify unique regions specific to target species. DNA target selection was successfully conducted using the comparative genomics tool from Genoscope, which has already proven to be effective in selecting DNA targets for developing PCR and qPCR tests for *Xanthomonas citri* pv. *citri* [48], as well as a LAMP test for the *R. pseudosolanacearum* phylotype I [37].

The BXW-LAMP assay designed in this study from a gene encoding for a histone-like nucleoid-structuring protein, was found to be 100% specific, both experimentally on the collection of target and non-target strains, and *in silico* on the 7,483 genomes analyzed. Nakato et al. [31] earlier established the exclusivity of this marker gene based on a 28 genomes database; we confirmed these results based on a much larger genome collection.

Conversely, the previously published BXW-specific LAMP [33] was found to be non-specific. Our extensive *in silico* analysis indeed identified a perfect match between the Hodgetts’ primers and the genomes of *X. vasicola* pv. *vasculorum* and *X. vasicola* pv. *zea* strain*s*, as well as a very high similarity score with other non-target species like *Xanthomonas cucurbitae* (96.9%) (data not shown).

The BBD-LAMP assay developed in this study, which amplified a DNA fragment in a putative hemolysin activation/secretion protein, proved to be highly specific, fully inclusive and exclusive both from *in silico* and molecular biology analyses. Specifically it outcompeted in inclusivity a visual LAMP assay recently published for the detection of *R. syzygii* subsp. *celebesensis* [34]. Indeed, the marker region of this visual LAMP assay was found absent in three of the eight genome sequences available on NCBI (data not shown), which could result in false negatives.

Since its first description in British Guiana in the 1850s [8] then Trinidad in the 1890s [49], Moko disease has been endemic in Central and South America, and is also present in the Southern Philippines, where the disease is known as Bugtok disease. The four sequevars IIB-3, IIB-4, IIA-6 and IIA-24 have been the most significant globally both in prevalence and virulence, regularly isolated from the 1950s until now, from various Musaceae in numerous countries across South and Central America, the Caribbean, and the Philippines (for IIB-3) [14,50–52]. These four sequevars were thus identified as priority targets for epidemiological monitoring.

The development of the duplex-MOKO LAMP assay faced different challenges: Moko strain genetic variability, genetic proximity of the banana-non-pathogenic “Emerging ecotype”(IIB-4NPB) [53]. The duplex-MOKO LAMP assay developed in this study detected 40 out of 41 strains tested, belonging to the four historical sequevars IIB-3, IIB-4, IIA-6 and IIA-24 by amplifying one or two DNA fragments, encoding for a conserved protein of unknown function (IIA) and for a conserved zinc ribbon domain-containing protein (IIB). Interestingly, the IIA marker was amplified from the sequevars IIA-6, IIA-24 and IIB-3, and the II-B marker from the IIB-4 and IIA-24 sequevars. These results suggest horizontal gene transfer between MOKO lineages, as previously mentioned before [53]. The only target strain not amplified by the duplex MOKO-LAMP assay was the IIB-3 strain RUN579 (also named UW11). This strain, along with the IIB-3 strains PD1446, UW75, and UW136, were isolated in the 1950s in Costa Rica. From the *in silico* study, none of these four strains contained the IIA LAMP marker in their genomes. Moreover, phylogenetic analysis based on 44,522 non-recombinant SNPs obtained from whole-genome alignment revealed that PD1446, UW75, UW136, and RUN579 clustered into a distinct subgroup within sequevar IIB-3 (S3 Fig.). Besides, the strain RUN579 was found to be non-pathogenic when inoculated on Cavendish. Together, these findings suggest that that RUN579 (together with the three-strain group) is actually an atypical Moko strain that requires further investigation, but it is not representative of the current Moko situation. Additionally, extended studies are also required on the two IIB-4 strains UA-1579 and UA-1609, described as pathogenic to banana [46] and yet unresponsive to the MOKO IIB target. We verified their genomes did not contain the MOKO IIB-4 target, but surprisingly they contained the IIB-4 NPB-specific amplicon [27] as well as other specific IIB-4 NPB markers described in [53].

None of the 91 non-target strains were amplified with the duplex MOKO-LAMP assay, except for a strain belonging to *R. thomasii*, isolated in a hospital setting, which tested positive although the fluorescence signal was delayed compared to that of an IIB-4 Moko strain. However, this strain was isolated from a human infection and is unlikely to be found on banana plants. The same applies to the 13 other genomes that matched *in silico* with the MOKO IIB primer set at a similar identity level (94%), all of which were isolated from different environments unrelated to banana tissues. Most importantly, the MOKO-LAMP assay did not detect any of the 54 non-target strains isolated from *Musa* spp. plants or the banana environment, particularly the IIB-4 NPB strains, which can be isolated from banana plants but are not pathogenic to this host.

Three sequevars have recently been described in Brazil within the Moko-inducing strains, namely IIB-25, IIA-41, and IIA-53 [16]. Only three genomes (2 for IIA-53 and 1 for IIB-25) are available in NCBI database. No match was found between the duplex MOKO primers and these three genomes. No marker common to these lineages could be found by [28] and two independent PCR assays (one duplex and one triplex PCR) were developed by the authors for detecting these particular Brazilian strains. As no strains are available in international collections, the development and validation of specific detection tests targeting these original groups is challenging. At present, the most appropriate approach would be to employ the RSSC-specific LAMP developed by [54]and refined by [37] as a field diagnostic tool in Brazil, complemented by the duplex MOKO-PCR, and, if necessary, followed by a more detailed analysis using the PCR methods described by [28].

High specificity can already be asserted for the duplex MOKO-LAMP test, regarding the most globally significant Moko sequevars, although the status of certain strains still requires verification. We would like to emphasize the relevance of conducting *in silico* studies in addition to experimental testing on bacterial strains to assess specificity. Indeed, our results showed that the experimental data was strongly supported by the *in silico* findings. *In silico* analyses provided a tremendous increase of the analytical power, allowing the screening of a large number of genomes for specificity, which would be very difficult to achieve through strain testing alone.

The development of the LAMP tests was accompanied by the elaboration of a nucleic acid extraction method suitable for field use. To achieve this, we utilized a NaOH-based method [37], that was improved by adding 2% polyvinylpyrrolidone (PVP) to neutralize potential plant compounds that inhibit the polymerase [55]. No statistical differences in the LAMP responses were observed when using this simplified extraction technique versus the extraction performed with a commercial kit on artificially infected plants, supporting the tolerance of the LAMP to plant inhibitors.

The sensitivity of LAMP assays was estimated between 0.1 pg/µl and 1 pg/µl (from 16 copies to 191 copies per reaction according to the different pathogens). These values are in accordance with the range found for other published LAMP assays [56–58].

When tested on tissues spiked with serially diluted bacteria, and extracted using the simplified extraction method, the LOD values ranged between 10^4^-10^5^ CFU/ml, corresponding to 50-500 copies per reaction. The LOD of 10^5^CFU/ml was obtained for three out of four Moko strains tested with the duplex MOKO-LAMP. Interestingly, the IIA-24 strain RUN20 displayed a LOD of 10^4^ CFU/ml, likely because both Moko markers were amplified, thereby generating more fluorescence., Regardless, a detection limit of 10^5^ CFU/ml, and *a fortiori* of 10^4^ CFU/ml, is more than sufficient to detect the pathogen in symptomatic material.

The LAMP assays considered in this study were validated on banana plants that were artificially inoculated with various bacterial strains in greenhouse conditions, as well as on samples collected from the field. Healthy samples consistently tested negative. For the detection from banana plants artificially inoculated in the greenhouse, consistent results were obtained between expression of internal/external symptoms, LAMP results and bacterial concentrations isolated from inoculated plants.

The field experiments, particularly in Indonesia, illustrated that symptomatology alone is insufficient in complex situations where multiple pathogens can cause similar symptoms.

The LAMP results obtained on field samples showed good concordance with the other PCR/qPCR methods targeting different DNA regions. Overall, LAMP results outperformed those of PCR or qPCR when these assays were conducted on the same simplified extracts. Besides, the absence of PCR or qPCR amplification in some LAMP-positive samples can most likely be attributed to the lower tolerance of PCR to plant inhibitors in a simplified extraction context, compared to LAMP.

The possibility of multiplexing several LAMP assays to simultaneously detect various pathogens could be explored. Some limitations include potential reduced sensitivity, and amplification of one pathogen masking the other. This was for example observed in our study with the duplex MOKO-LAMP, where the IIB-specific marker was preferentially amplified in IIA-24 strains. Despite these challenges, multiplexing could be particularly useful for pathogens with similar symptoms that co-occur in the same areas, such as Moko and Fusarium wilt which challenges surveillance effort.

## Conclusion

Effective surveillance, supported by accurate diagnostic tools, is essential to prevent pathogen introduction and spread. Moreover, preventing the introduction and establishment of major pests and diseases is widely recognized as being far more cost-effective than implementing control or eradication measures once they are established. The LAMP assays developed in this project are specifically optimized for on-site application, unlike existing molecular diagnostic methods that require a laboratory setting. The developed detection assay can be integrated into diagnostic programs aimed at protecting borders against these bacterial pathogens that pose a threat to global food security and agricultural economies.

## Acknowledgments

We are very grateful to Lorène Milliex and Tetiana Kwan-Tat for their involvement in the follow-up of the project, Caroline Chatillon and Camilo Gianinazzi for their technical support.

## Supporting information

**S1 Fig. Areas where the field surveys were conducted**

**(A)** Uganda, (**B**) Indonesia (Java), (**C**) Ecuador, and (**D**) Jamaica. The numbers of samples collected and tested by LAMP are indicated in red.

**S2 Fig. Percentage of diseased plants over time following inoculation with the different pathogens**

Tendency curves with their 95% confidence intervals were calculated using local polynomial regressions (locally estimated scatterplot smoothing [LOESS] method).

**S3 Fig. Phylogenetic tree of *Ralstonia solanacearum* sequevar IIB-3 strains**, using a sequevar IIB-25 strain as outgroup. The tree was constructed with RAxML as implemented in Gubbins v3.3.3, based on 44,522 nonrecombinant single nucleotide polymorphisms (SNPs) obtained from whole-genome alignment using Parsnp v1.7.4. Branch support values were calculated from 1,000 bootstrap replicates. The tree scale indicates the number of substitutions per genome.

**S1 Table. BXW and non target strains used in this study for BXW-LAMP assay specificity. S2 Table. BBD and non target strains used in this study for BBD-LAMP assay specificity.**

**S3 Table. Moko and non target strains used in this study for MOKO duplex-LAMP specificity.**

**S4 Table. Target and non target genomes used for diagnostic markers selection**

The selection was performed using the “gene phyloprofile “ tool of the Microscope plateform (Genoscope, France). The strains corresponding to genomes publicly available on the platform (https://mage.genoscope.cns.fr/microscope/home/index.php) are bolded . Genome accession found on NCBI were also added.

**S5 Table. Selected DNA targets**

**S6 Table. Genomes used for the *in silico* analyse and % identity between the BXW-specific target located in a gene coding for a histone-like nucleoid-structuring protein, and the corresponding LAMP primers sets.**

**S7 Table. Genomes used for the *in silico* analyse and % identity between the BBD-specific target located in a gene coding for a putative hemolysin activation/secretion protein, and the corresponding LAMP primers sets.**

**S8 Table. Genomes used for the *in silico* analyse and % identity between the MOKO-specific targets located in a gene coding for a conserved protein of unknown function (IIA) and a gene coding for a conserved zinc ribbon domain-containing protein (IIB) and the corresponding LAMP primers sets.**

## References

1. Bebber DP. Range-Expanding Pests and Pathogens in a Warming World. Annu Rev Phytopathol. 2015 Aug 4;53(1):335–56.

2. Miller SA, Beed FD, Harmon CL. Plant disease diagnostic capabilities and networks. Annu Rev Phytopathol. 2009 Sept 1;47(1):15–38.

3. FAO. FAOSTAT. 2022.

4. Blomme G, Dita M, Jacobsen KS, Pérez Vicente L, Molina A, Ocimati W, et al. Bacterial Diseases of Bananas and Enset: Current State of Knowledge and Integrated Approaches Toward Sustainable Management. Front Plant Sci. 2017 July 20;8:1290.

5. Prior P, Ailloud F, Dalsing BL, Remenant B, Sanchez B, Allen C. Genomic and proteomic evidence supporting the division of the plant pathogen *Ralstonia solanacearum* into three species. BMC Genomics. 2016 Dec;17(1):90.

6. Safni I, Cleenwerck I, De Vos P, Fegan M, Sly L, Kappler U. Polyphasic taxonomic revision of the *Ralstonia solanacearum* species complex: proposal to emend the descriptions of *Ralstonia solanacearum* and *Ralstonia syzygii* and reclassify current *R. syzygii* strains as *Ralstonia syzygii* subsp. *syzygii* subsp. nov., *R. solanacearum* phylotype IV strains as *Ralstonia syzygii* subsp. *indonesiensis* subsp. nov., banana blood disease bacterium strains as *Ralstonia syzygii* subsp. *celebesensis* subsp. nov. and *R. solanacearum* phylotype I and III strains as *Ralstonia pseudosolanacearum* sp. nov. Int J Syst Evol Microbiol. 2014 Sept 1;64(Pt_9):3087–103.

7. Álvarez, Elizabeth, Pantoja, Alberto, Gañan, Lederson, Ceballos, Germán. Current status of Moko disease and black sigatoka in Latin America and the Caribbean, and options for managing them. Centro Internacional de Agricultura Tropical (CIAT), Cali, CO. 50 p. (Publicación CIAT No. 404); 2015.

8. Sequeira L. Bacterial Wilt: the Missing Element in International Banana Improvement Programs. In: Prior P, Allen C, Elphinstone J, editors. Bacterial Wilt Disease [Internet]. Berlin, Heidelberg: Springer Berlin Heidelberg; 1998 [cited 2025 Oct 15]. p. 6–14. Available from: http://link.springer.com/10.1007/978-3-662-03592-4_2

9. Zulperi D, Sijam K. First Report of *Ralstonia solanacearum* Race 2 Biovar 1 Causing Moko Disease of Banana in Malaysia. Plant Disease. 2014 Feb;98(2):275–275.

10. Belalcazar, S. C, Rosales, F. E, Pocasangre, L. E. El Moko del banano y el plátano y el rol de las plantas hospederas en su epidemiología. In: Proceedings of the XVI International ACORBAT Meeting. Oaxaca, Mexico: eds M. Orozco-Santos, J. Orozco-Romero, M. Robles-Gonzalez, J. Velazquez-Monreal, V. Medina-Urrutia, and J. A. Hernandez-Bautista (Oaxaca: Artturi); 2004. p. 16–35.

11. Delgado R, Morillo E, Buitrón J, Bustamante A, Sotomayor I. First report of Moko disease caused by *Ralstonia solanacearum* race 2 in plantain (*Musa* AAB) in Ecuador. New Disease Reports. 2014 July;30(1):23–23.

12. Redacción El Universo. Enfermedad del moko afecta a 2.700 hectáreas de banano en Los Ríos, según resultados de proyecto de AEBE. El Universo [Internet]. 2025 Aug 30; Available from: https://www.eluniverso.com/noticias/economia/enfermedad-del-moko-afecta-a-2700-hectareas-de-banano-en-los-rios-segun-resultados-de-proyecto-de-aebe-nota/

13. Corazón, Vera. Boletín Técnico N° 07-2025-IPC [Internet]. Ecuador: El Instituto Nacional de Estadística y Censos (INEC); 2025 July. Report No.: N° 07-2025-IPC. Available from: https://www.ecuadorencifras.gob.ec/documentos/web-inec/Inflacion/2025/Julio/Boletin_tecnico_07-2025-IPC.pdf

14. Fegan M, Prior P. Diverse members of the *Ralstonia solanacearum* species complex cause bacterial wilts of banana. Austral Plant Pathol. 2006;35(2):93.

15. Wicker E, Grassart L, Coranson-Beaudu R, Mian D, Guilbaud C, Fegan M, et al. *Ralstonia solanacearum* Strains from Martinique (French West Indies) Exhibiting a New Pathogenic Potential. Appl Environ Microbiol. 2007 Nov;73(21):6790–801.

16. Albuquerque GMR, Santos LA, Felix KCS, Rollemberg CL, Silva AMF, Souza EB, et al. Moko Disease-Causing Strains of *Ralstonia solanacearum* from Brazil Extend Known Diversity in Paraphyletic Phylotype II. Phytopathology®. 2014 Nov;104(11):1175–82.

17. Buddenhagen I. BLOOD BACTERIAL WILT OF BANANA: HISTORY, FIELD BIOLOGY AND SOLUTION. Acta Hortic. 2009 May;(828):57–68.

18. Teng SK, Abd. Aziz NA, Mustafa M, Laboh R, Ismail IS, Rohani Sulaiman S, et al. The Occurrence of Blood Disease of Banana in Selangor, Malaysia. IJAB. 2015 Oct 1;18(01):92–7.

19. Safni I, Subandiyah S, Fegan M. Ecology, Epidemiology and Disease Management of Ralstonia syzygii in Indonesia. Front Microbiol. 2018 Mar 13;9:419.

20. Thwaites, Mansfield, Eden-Green, Seal. RAPD and rep PCR-based fingerprinting of vascular bacterial pathogens of *Musa* spp. Plant Pathology. 1999 Feb;48(1):121–8.

21. Nakato V, Mahuku G, Coutinho T. *Xanthomonas campestris* pv. *musacearum* : a major constraint to banana, plantain and enset production in central and east Africa over the past decade. Molecular Plant Pathology. 2018 Mar;19(3):525–36.

22. Geberewold AZ. Review on impact of banana bacterial wilt (*Xanthomonas campestris pv. Musacerum*) in East and Central Africa. Yildiz F, editor. Cogent Food & Agriculture. 2019 Jan 1;5(1):1586075.

23. Nakato GV, Studholme DJ, Blomme G, Grant M, Coutinho TA, Were EM, et al. SNP-based genotyping and whole-genome sequencing reveal previously unknown genetic diversity in *Xanthomonas vasicola* pv. *musacearum*, causal agent of banana xanthomonas wilt, in its presumed Ethiopian origin. Plant Pathology. 2021 Apr;70(3):534–43.

24. Nakato GV, Fuentes Rojas JL, Verniere C, Blondin L, Coutinho T, Mahuku G, et al. A new Multi Locus Variable Number of Tandem Repeat Analysis Scheme for epidemiological surveillance of Xanthomonas vasicola pv. musacearum, the plant pathogen causing bacterial wilt on banana and enset. Chiang TY, editor. PLoS ONE. 2019 Apr 11;14(4):e0215090.

25. PHA. Bacterial Wilt of Banana Diagnostics Manual., S. Porchum, ed. Cooperative Research Centre For Tropical Plant Protection, Plant Health Australia.; 2006.

26. Prior P, Fegan M. Diversity and molecular detection of *Ralstonia solanacearum* race 2 strains by multiplex PCR. In: Allen C, Prior P, Hayward AC, editors. Bacterial wilt disease and the Ralstonia solanacearum species complex. St. Paul, Minn: American Phytopathological Society; 2005. p. 405–14.

27. Cellier G, Moreau A, Chabirand A, Hostachy B, Ailloud F, Prior P. A duplex PCR assay for the detection of *Ralstonia solanacearum* Phylotype II strains in *Musa* spp. Lee SW, editor. PLoS ONE. 2015 Mar 26;10(3):e0122182.

28. Rincón-Flórez VA, Carvalhais LC, Silva AMF, McTaggart A, Ray JD, O’Dwyer C, et al. Validation of PCR Diagnostic Assays for Detection and Identification of All *Ralstonia solanacearum* Sequevars Causing Moko Disease in Banana. Phytopathology®. 2024 Nov 1;114(11):2375–84.

29. Das, S. Molecular diagnostics of the Blood Disease Bacterium. Honors. Thesis. University of Queensland, Brisbane, Australia. Davis, R., Fegan,. [Brisbane, Australia.]: Honors Thesis. University of Queensland; 2004.

30. Rincón-Flórez VA, Ray JD, Carvalhais LC, O’Dwyer CA, Subandiyah S, Zulperi D, et al. Diagnostics of Banana Blood Disease. Plant Disease. 2022 Mar 1;106(3):947–59.

31. Nakato GV, Wicker E, Coutinho TA, Mahuku G, Studholme DJ. A highly specific tool for identification of *Xanthomonas vasicola* pv. *musacearum* based on five Xvm-specific coding sequences. Heliyon. 2018;4(12):e01080.

32. Notomi T. Loop-mediated isothermal amplification of DNA. Nucleic Acids Res. 2000 June 15;28(12):63e–63.

33. Hodgetts J, Hall J, Karamura G, Grant M, Studholme DJ, Boonham N, et al. Rapid, specific, simple, in-field detection of *Xanthomonas campestris* pathovar *musacearum* by loop-mediated isothermal amplification. J Appl Microbiol. 2015 Dec;119(6):1651–8.

34. Azizi MMF, Lau HY, Abu Bakar N, Romeli S, Mohd Yusof MF, Badrun R, et al. Development of a Highly Sensitive Loop-Mediated Isothermal Amplification Incorporated with Flocculation of Carbon Particles for Rapid On-Site Diagnosis of Blood Disease Bacterium Banana. Horticulturae. 2022 May 5;8(5):406.

35. Kelman A. The Relationship of Pathogenicity of Pseudomonas solanacearum to Colony Appearance in a Tetrazolium Medium. Phytopathology. 1954;693–5.

36. Vallenet D, Calteau A, Dubois M, Amours P, Bazin A, Beuvin M, et al. MicroScope: an integrated platform for the annotation and exploration of microbial gene functions through genomic, pangenomic and metabolic comparative analysis. Nucleic Acids Research. 2019 Oct 24;gkz926.

37. Robène I, Maillot-Lebon V, Pecrix Y, Ravelomanantsoa S, Massé D, Jouen E, et al. Loop-Mediated Isothermal Amplification Assays for Rapid Detection of *Ralstonia solanacearum* Species Complex and Phylotype I in Solanaceous Crops. PhytoFrontiersTM. 2023 Oct 30;PHYTOFR-08-22-0089-R.

38. Anonymous. PM 7/98 (5) Specific requirements for laboratories preparing accreditation for a plant pest diagnostic activity. EPPO Bulletin. 2021 Dec;51(3):468–98.

39. Winstead, N., Kelman, A. Inoculation Techniques for Evaluating Resistance to Pseudomonas solanacearum. Phytopathology. 1952;628–34.

40. Ahmadi E, Kowsari M, Azadfar D, Salehi Jouzani G. Rapid and economical protocols for genomic and metagenomic DNA extraction from oak (Quercus brantii Lindl.). Annals of Forest Science. 2018 June;75(2):43.

41. Elphinstone JG, Hennessy J, Wilson JK, Stead DE. Sensitivity of different methods for the detection of Ralstonia solanacearum in potato tuber extracts. EPPO Bulletin. 1996 Sept;26(3–4):663–78.

42. Cenis JL. Rapid extraction of fungal DNA for PCR amplification. Nucl Acids Res. 1992;20(9):2380–2380.

43. Gandelman O, Jackson R, Kiddle G, Tisi L. Loop-mediated amplification accelerated by stem primers. Int J Mol Sci. 2011 Dec 8;12(12):9108–24.

44. Beutler J, Holden S, Georgoulis S, Williams D, Norman DJ, Lowe-Power TM. Whole Genome Sequencing Suggests that “Nonpathogenicity on Banana (NPB)” Is the Ancestral State of the *Ralstonia solanacearum* IIB-4 Lineage. PhytoFrontiers^TM^. 2023 Sept;3(2):262–7.

45. Pais AKL, Santos LVSD, Albuquerque GMR, Farias ARGD, Silva Junior WJ, Balbino VDQ, et al. Comparative genomics and phylogenomics of the *Ralstonia solanacearum* Moko ecotype and its symptomatological variants. Genet Mol Biol. 2022;45(4):e20220038.

46. Ramírez M, Moncada RN, Villegas-Escobar V, Jackson RW, Ramírez CA. Phylogenetic and pathogenic variability of strains of *Ralstonia solanacearum* causing moko disease in Colombia. Plant Pathology. 2020 Feb;69(2):360–9.

47. Moehling TJ, Browne ER, Meagher RJ. Effects of single and multiple nucleotide mutations on loop-mediated isothermal amplification. Analyst. 2024;149(6):1701–8.

48. Robène I, Maillot-Lebon V, Chabirand A, Moreau A, Becker N, Moumène A, et al. Development and comparative validation of genomic-driven PCR-based assays to detect *Xanthomonas citri* pv. *citri* in citrus plants. BMC Microbiol. 2020 Dec;20(1):296.

49. Rorer JB. A bacterial disease of bananas and plantains. Phytopathology. 1911;45–9.

50. Fegan M. Bacterial wilt diseases of banana: evolution and ecology. In: Bacterial wilt disease and the Ralstonia solanacearum species complex. C. Allen, P. Prior, A. C. Hayward. Saint-Paul USA: American Phytopathological Society; 2005. p. 379–86.

51. Lowe-Power TM, Avalos J, Bai Y, Munoz MC, Chipman K, Elmgreen VN, et al. A Meta-analysis of the known Global Distribution and Host Range of the *Ralstonia* Species Complex [Internet]. 2020 [cited 2025 Mar 13]. Available from: http://biorxiv.org/lookup/doi/10.1101/2020.07.13.189936

52. Sharma P, Johnson MA, Mazloom R, Allen C, Heath LS, Lowe-Power TM, et al. Meta-analysis of the *Ralstonia solanacearum* species complex (RSSC) based on comparative evolutionary genomics and reverse ecology. Microbial Genomics [Internet]. 2022 Mar 1 [cited 2025 May 20];8(3). Available from: https://www.microbiologyresearch.org/content/journal/mgen/10.1099/mgen.0.000791

53. Ailloud F, Lowe T, Cellier G, Roche D, Allen C, Prior P. Comparative genomic analysis of *Ralstonia solanacearum* reveals candidate genes for host specificity. BMC Genomics. 2015 Dec;16(1):270.

54. Okiro LA, Tancos MA, Nyanjom SG, Smart CD, Parker ML. Comparative evaluation of LAMP, qPCR, conventional PCR, and ELISA to detect *Ralstonia solanacearum* in Kenyan potato fields. Plant Dis. 2019 May;103(5):959–65.

55. Rezadoost MH, Kordrostami M, Kumleh HH. An efficient protocol for isolation of inhibitor-free nucleic acids even from recalcitrant plants. 3 Biotech. 2016 June;6(1):61.

56. Ordóñez N, Salacinas M, Mendes O, Seidl MF, Meijer HJG, Schoen CD, et al. A loop-mediated isothermal amplification (LAMP) assay based on unique markers derived from genotyping by sequencing data for rapid *in planta* diagnosis of Panama disease caused by Tropical Race 4 in banana. Plant Pathology. 2019 Dec;68(9):1682–93.

57. Patel R, Mitra B, Vinchurkar M, Adami A, Patkar R, Giacomozzi F, et al. A review of recent advances in plant-pathogen detection systems. Heliyon. 2022 Dec;8(12):e11855.

58. Venbrux M, Crauwels S, Rediers H. Current and emerging trends in techniques for plant pathogen detection. Front Plant Sci. 2023 May 8;14:1120968.

